# Physiological cerebrospinal fluid like medium reveals autophagy dependency of leukaemia in the central nervous system

**DOI:** 10.64898/2026.03.09.709824

**Authors:** Ekaterini Himonas, Anand Manoharan, Kiron Roy, Kevin M. Rattigan, Angela Ianniciello, Martha M. Zarou, Daniele Sarnello, Leenus Martin, Robert Shoemaker, David Sumpton, Saverio Tardito, Chris Halsey, G. Vignir Helgason

**Author notes:** **Corresponding authors**: &. Joint senior authors (Chris Halsey, G. Vignir Helgason).

## Abstract

Nutrient availability is a critical environmental factor that influences the metabolism and adaptability of cancer cells, including acute lymphoblastic leukaemia (ALL) cells, prone to relapse in the central nervous system (CNS). Currently available cell culture media contain supraphysiological nutrient levels and do not represent the restricted metabolic environment of CNS-ALL which resides in the leptomeninges surrounded by cerebrospinal fluid (CSF). Therefore, we formulated a novel physiological CSF-like cell culture medium (CSFmax) that recapitulates the unique metabolite composition of the CSF. Through *in vitro* and *in vivo* metabolic and functional studies, we demonstrate that ALL cells cultured in CSFmax rewire their metabolism, closely mimicking the metabolic phenotype of CNS-ALL, including their metabolic activity and redox state. Utilising CSFmax, in comparison to conventional nutrient-rich culture media, we identified an essential role for autophagy in ALL adaptation to the CNS niche. This was evident by increased autophagic activity and selective sensitisation of ALL cells to pharmacological inhibition of autophagy and genetic knockout of Unc-51 Like Autophagy Activating Kinase 1 (ULK1) or autophagy related 7 (ATG7). Importantly, using a robust preclinical *in vivo* model, mice xenografted with ULK1 and ATG7 deficient ALL cells exhibited reduced CNS disease burden when compared to mice xenografted with control cells. Overall, our findings provide strong evidence that physiological CSFmax is superior to current *in vitro* culture systems in recapitulating the metabolic signature of CNS resident ALL cells. By exploiting this system, we revealed for the first time autophagy as a targetable therapeutic vulnerability in CNS-ALL.

**Key Points:**

- Culturing ALL cells in bespoke CSF-like medium (CSFmax) recapitulates the metabolic adaptation of ALL cells in the CNS niche
- Autophagy is critical for metabolic adaptation and survival of CNS resident ALL cells

## Introduction

Nonphysiological concentrations of various nutrients in commercial cell culture media may rewire cancer cellular metabolism and generate undesired artificial phenotypes [1–4]. This led to the recent development of more physiological cell culture media that mimic the physiology of human plasma and more closely recapitulate the metabolic phenotype of cells *in vivo* [5, 6]. However, currently there is no commercial medium available that mimics distinct metabolic microenvironments, such as the cerebrospinal fluid (CSF).

CSF surrounds the brain and spinal cord within the leptomeninges. It is a mixture of water, ions, nutrients, and macromolecules which is frequently replenished to allow the removal of waste products from the central nervous system (CNS) [7, 8]. Therefore, CSF ensures CNS homeostasis, facilitates communication between the CNS and other systems, and functionally supports cell populations inhabiting this niche by providing them energy and metabolic building blocks [9, 10]. Overall, the level of nutrients and oxygen (O_2_) are restricted in the CSF microenvironment compared to human plasma [7, 11–13]. Consequently, a cell culture medium that recapitulates the metabolite composition of CSF would be pivotal, particularly in studies related to leptomeningeal-tropic malignancies, such as acute lymphoblastic leukaemia (ALL).

ALL cells commonly migrate from the bone marrow (BM) to the CNS where they engraft within the leptomeninges surrounded by CSF, evading immune responses and systemic drug treatment [14–17]. As such, optimal eradication of CNS-ALL remains one of the major challenges in treatment of childhood ALL [18–23]. We have shown that it is a generic, non-clonal property of ALL cells to cross the blood-CSF barrier [24], and subsequent metabolic alterations are required to adapt to the nutrient-deprived and hypoxic CNS microenvironment [25–27]. However, further investigation is required to fully elucidate and exploit the adaptive metabolic rewiring mechanisms ALL cells undergo in the CNS niche.

Therefore, we formulated a physiological CSF-like cell culture medium, herein called CSFmax, comprising nutrients and metabolites at the concentration found in human CSF. Through metabolic characterisation of ALL cells cultured in CSFmax in 21% O_2_ (“normoxia”) and in 1% O_2_ levels (“hypoxia”), in comparison to relatively nutrient-rich conventional media, we demonstrate that the metabolic functions of cells cultured in CSFmax are comparable to leukaemic cells freshly isolated from CNS, with a reduction in their overall energy status in conjunction with greater oxidative stress. Importantly, by utilising CSFmax and robust pre-clinical xenograft models we identify an essential role for autophagy in the adaptation of ALL cells to CSF conditions. Overall, we demonstrate the potential of CSFmax to uncover fundamental adaptive mechanisms to the nutrient-scarce CSF niche, applicable to CNS leukaemia, other leptomeningeal-tropic malignancies, and CNS autoimmune disorders.

## Methods

### Formulation of CSF-like medium (CSFmax)

CSFmax is a novel physiological medium developed to closely mimic the key nutrient composition of human CSF. Nutrient and metabolite concentrations of the CSF from healthy individuals were estimated from the values reported in the Human Metabolome database [12] from which the mean concentrations were adopted. The concentration of all components of CSFmax are listed in Supplemental Data.

### Cell culture

ALL cell lines were cultured in RPMI 1640 medium (Cat. #11875093) supplemented with 100 IU/mL penicillin/streptomycin (Cat. #15140122), 2 mM L-glutamine (Cat. #A2916801), and 10% (v/v) fetal bovine serum (FBS, Cat. #10100147). ALL cell lines were also cultured in Plasmax^TM^ (Cat. #156371) [6] supplemented with 100 IU/mL penicillin/streptomycin (Cat. #15140122) and 10% (v/v) dialysed fetal bovine serum (FBS, Cat. #A3382001). Finally, ALL cell lines were also cultured in CSFmax formulated as described above (see section Formulation of CSF-like medium (CSFmax)).

Cell lines were maintained at 5% CO_2_ and 37 °C. Cell number and viability were monitored using a CASY automated cell counter (Roche). All cell lines were obtained from DSMZ and were authenticated and regularly tested for mycoplasma contamination. Where indicated, the following compounds were added to the medium: 50 nM bafilomycin A1 (Cat. #A8627-APE), 1 μM MRT403 (Erasca), 100 nM cytarabine (Cat. #PHR1787).

### Generation of KO and stable cell lines

To generate CRISPR-Cas9-mediated gene knockouts of BNIP3, ULK1, and ATG7 in ALL cell lines, we used guide sequences as reported (Supplementary Table 3). Synthesis of selected guides was conducted by Integrated DNA Technologies. Oligonucleotides were annealed and cloned in BsmBl-digested lentiCRISPR v2 plasmid containing a puromycin-resistance marker (Cat. #52961). Lentiviral particles for cloned lentiCRISPR v2 were generated by transfecting human embryonic kidney (HEK) 293FT cells with pCMV-VSV-G envelope and psPAX2 packaging vectors using the calcium phosphate method [28]. Viral supernatant was harvested after 48 h and transferred onto target ALL cells. Cells were then selected using 3 μg/mL puromycin (Cat. #12122530) and presence of guides was validated by immunoblotting.

Similarly, to generate stable ALL cell lines expressing the fluorescent probe GFP-LC3-RFP-LC3ΔG we used PMRX-IP-GFP-LC3-RFP which was a gift from Noboru Mizushima (Addgene Plasmid #84573) [29]. Cells were then selected using 3 μg/mL puromycin (Cat. #12122530) and were isolated by sorting using FACSAria Fusion Cell sorter (BD Biosciences).

### Animal studies

The NOD/SCID-IL2Rγ^−/−^, NOD.Cg-Prkdc^scid^Il2rγ^tm1Wjl^/SzJ (NSG) mice strain were purchased from Charles River Laboratories, Harlow, United Kingdom. Animals were housed in a pathogen-free facility at the Cancer Research UK Scotland Institute and kept in day/night cycles (12 h each), with temperature of 20–24 °C and humidity 45–65%. Feeding and water were ad libitum. Mice weights and general health scores were monitored daily until they reached experimental endpoint due to observed clinical signs according to institutional guidelines. Overall, mice transplanted with SEM vector control cells reached experimental endpoint on day 21 as they reached the overall severity threshold.

### Cell line xenograft experiments

Genetically interfered cell lines were produced as described above (see section Generation of KO stable cell lines). SEM vector control, BNIP3 knockout (KO), ULK1 KO, or ATG7 KO cells (2×10^6^ cells per mouse) were transplanted via tail vein injection into female and male NSG mice aged 6-12 weeks. All mice from all experimental arms were sacrificed on day 21 and brain, BM from the hip, tibia, and femur, as well as spleen were harvested from each mouse.

### Study approval

Animal experiments were performed according to all applicable laws and regulations with ethical approval from the University of Glasgow Animal Welfare and Ethical Review Board (AWERB) under approved project licence PPL PP5432144 and personal licence PIL I98283172.

### Statistics

Statistical analyses were performed using Prism 9 GraphPad Software. The number of biological replicates and statistical tests employed are described in figure legends. Data are presented as mean ± s.e.m as indicated. Results were considered statistically significant when p values ≤ 0.05. For *in vitro* experiments, no statistical method was used to predetermine sample size; however, at least 3 biological or 3 independent culture replicates were used to perform any statistical analysis. Cells were randomly plated/treated/analysed during each experiment for all *in vitro* experiments. Investigators were not blinded during data processing and analysis of *in vitro* data. Statistical significance for parametric data was determined by paired or unpaired two-tailed t-tests. To assess statistical significance for three or more groups, one-way analysis of variance (ANOVA) was employed, while two-way ANOVA was used to determine significance between multiple groups with two experimental parameters. For *in vivo* experiments, similarly aged animals were transplanted with the same number of donor cells. Mice were randomly allocated to control, or experimental conditions and no animal was excluded from the analysis. Investigators were not blinded during data processing and analysis of *in vivo* data.

Further information is provided in the Supplemental Materials and Methods.

## Results

### CSFmax culture recapitulates the physiology of ALL cells in the CNS microenvironment

Currently available commercial cell culture media, including RPMI 1640 (RPMI) and Plasmax™ (Plasmax), are composed with glucose, amino acids, vitamins, and salts at concentrations that exceed those of human CSF (Supplementary Table 1). Therefore, we aimed to formulate and characterise a novel medium intended to closely mimic the metabolic environment of human CSF where ALL cells promote CNS relapse [14–17]. Previously reported concentrations of metabolites detected in CSF samples from healthy individuals were used to implement the culture medium, herein called CSFmax [12] (Figure 1A, Supplementary Table 1). Direct comparison of the formulation of CSFmax, with RPMI, Plasmax, and human CSF metabolome as reported, demonstrated that CSFmax closely recapitulates both the total composition of key nutrients and the individual concentration of indicated nutrients found in human CSF (Figure 1A-B, Supplementary Table 1). Besides key nutrients present in human CSF, other factors encompassing the CSF microenvironment include the lower levels of lipids and growth factors [11, 13] which play a critical role in cell proliferation. Therefore, in our experiments RPMI and Plasmax were supplemented with 10% fetal bovine serum (FBS) and 10% dialysed FBS, respectively, to model growth factor- and lipid-rich conditions, such as those encountered in BM. In contrast, CSFmax was formulated to provide comparatively lipid and growth factor-limited conditions and was therefore supplemented with 5% delipidated dialysed FBS (Supplementary Table 2).

**Figure 1:**
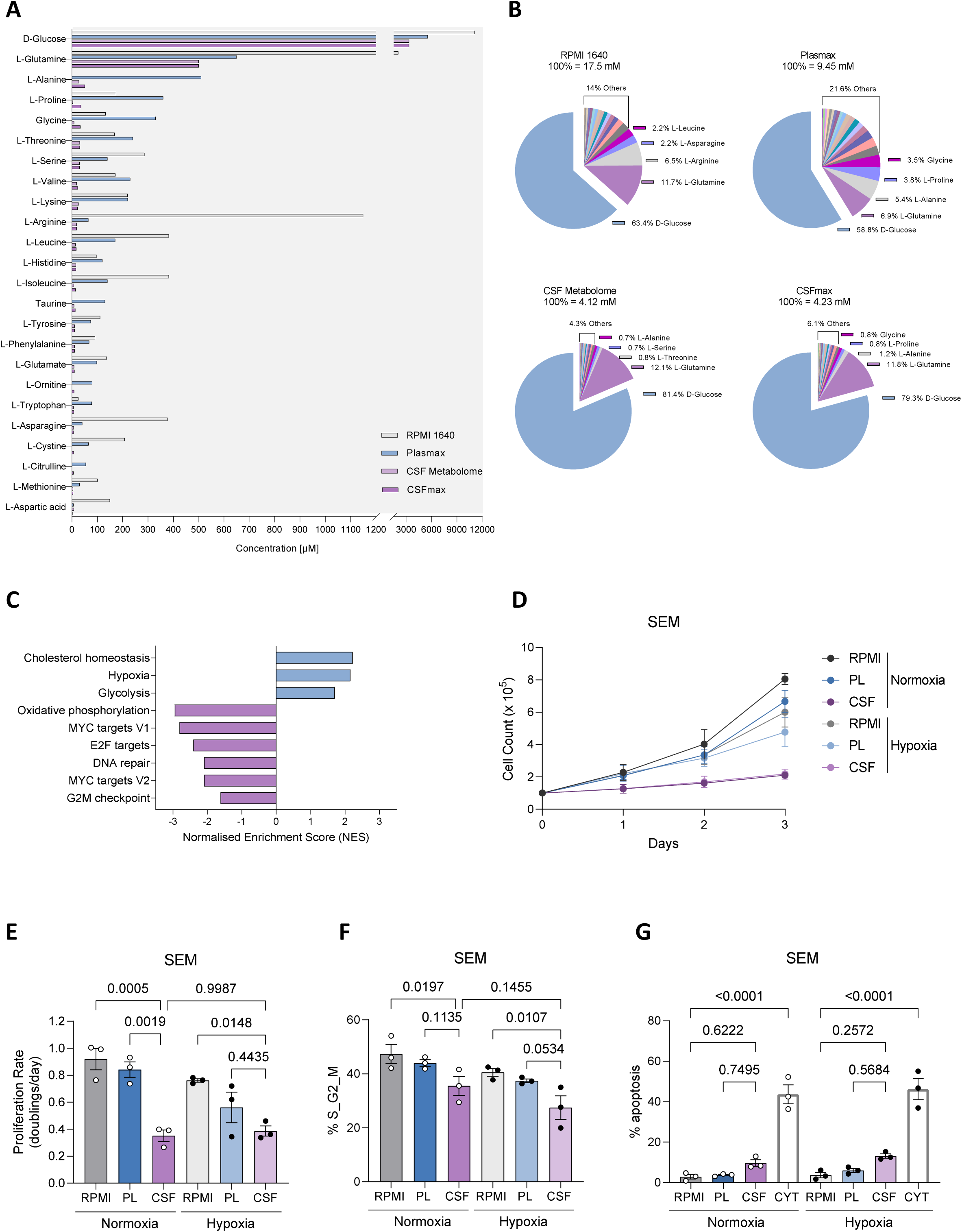
A novel cell culture medium that recapitulates the physiology of ALL cells in the CNS microenvironment. **A,** Direct comparison of the concentration of indicated metabolites composing CSFmax with RPMI, Plasmax, and the average CSF metabolome. See Supplementary Table 1 for further detail regarding the concentration of indicated metabolites. **B,** Total nutrient composition of RPMI, Plasmax, average CSF metabolome, and CSFmax of indicated metabolites in response to (A). **C,** Gene set enrichment analysis (GSEA) of significantly differentially expressed genes in human SEM and REH cells isolated from the CNS compared to human SEM and REH cells isolated from the spleen of xenografted mice (n=2 independent groups of 5 mice for SEM cells, n=2-3 independent groups of 5 mice for REH cells). Publicly available dataset used: GSE135115. GSEA hallmarks represent statistically significant biological functions upregulated and downregulated in human ALL cells isolated from the CNS. Blue bars indicate signatures with positive normalised enrichment score (NES) in CNS versus spleen samples, and purple bars indicate signatures with negative NES in CNS versus spleen samples. **D,E,** Growth curves (D) and proliferation rate (D) of SEM cells cultured in RPMI, Plasmax (PL), and CSFmax (CSF) in normoxia (N, 21% O_2_) or hypoxia (H, 1% O_2_) (n=3 independent experiments). **F,** Total percentage of S and G2_M cell cycle stages in SEM cells in response to culture conditions as indicated in (D, E) (n=3 independent experiments). **G,** Percentage of apoptosis in SEM cells in response to culture conditions as indicated in (D, E) for 72 h and treatment with 150 nM cytarabine (CYT) in RPMI for 72 h (n=3 independent experiments). Data are shown as the mean ± s.e.m. P-values were calculated with a repeated measure one-way ANOVA with Tukey’s multiple comparison test (E, G) and two-way ANOVA with Tukey’s multiple comparison test (F).

The BM is favoured by its cellular complexity and its dense vascularisation [30, 31], hence it accounts for higher levels of nutrients and growth factors compared to CSF. Firstly, we performed bioinformatic analysis where we assessed the transcriptional differences in human ALL cells isolated from the CNS compared to human ALL cells isolated from the periphery (spleen) of xenografted mice. While cholesterol homeostasis, hypoxia response and glycolysis were upregulated, gene set enrichment analysis (GSEA) revealed oxidative phosphorylation (OXPHOS) signature to be downregulated in CNS-derived leukaemic cells, as well as MYC targets, E2F targets, DNA repair, and G2/M checkpoint hallmarks (Figure 1C). Downregulation of these processes, which are involved in cell proliferation suggested that despite potential switch from mitochondrial metabolism to glycolysis, CNS-derived leukaemic cells are metabolically less active compared to leukaemic cells derived from the periphery. Indeed, we showed that SEM and REH cells cultured in CSFmax exhibited a significant decrease in growth and proliferation rate when compared to cells cultured in a conventional (RPMI) or BM-mimicking (Plasmax) cell culture medium [32] (Figure 1D-E, Supplementary Figure 1A-B). Notably, SEM and REH cells cultured in CSFmax in hypoxia did not exhibit a further reduction in growth nor proliferation rate, suggesting that the proliferation in CSF is mainly constrained by nutrient availability (Figure 1D-E, Supplementary Figure 1A-B). Importantly, SEM and REH cells cultured in CSFmax in either normoxia or hypoxia, were characterised by a reduction of cells entering the S/G2/M phase (Figure 1F, Supplementary Figure 1C-D), with a minimal increase in apoptotic cell numbers (Figure 1G, Supplementary Figure 1E). This is in line with previous work demonstrating that ALL cells retrieved from the CNS reduce their cell proliferation and undergo cell cycle arrest, when compared to ALL cells retrieved from the BM [25].

### CSFmax culture extensively alters the metabolic activity of ALL cells comparably to CNS-derived ALL cells

We next sought to explore alterations in metabolic activity in ALL cells cultured in CSFmax. Initially we wanted to exclude the possibility that the observed antiproliferative phenotype was exhibited due to nutrient exhaustion in the spent medium. Assessing the extracellular levels of metabolites at various timepoints, we observed that SEM cells cultured in CSFmax have key metabolites such as glucose, available to consume up to 72 hours (h), whilst SEM cells cultured in Plasmax consumed more than 80% of the available glucose during this period (Supplementary Figure 2A-B). This was in line with the significant reduction in the consumption rate of glucose in SEM cells cultured in CSFmax at 48 and 72 h of incubation, when compared to SEM cells cultured in Plasmax (Supplementary Figure 2B). Importantly, extracellular levels of lactate, a metabolite secreted during glucose oxidation, were modestly reduced in SEM cells cultured in CSFmax at 72 h of incubation, together with increased lactate consumption (Supplementary Figure 2C-D). By contrast, SEM cells cultured in Plasmax showed increased extracellular levels of lactate, with reduced levels of lactate consumption, suggesting increased lactate secretion (Supplementary Figure 2C-D). Surprisingly, the reduction in metabolite consumption in SEM cells cultured in CSFmax was not limited to glucose, as results indicated that over time, the consumption of almost all metabolites was significantly reduced when compared to SEM cells cultured in Plasmax (Figure 2A, Supplementary Figure 2E-F), further highlighting their reduced metabolic activity in CSFmax. Conversely, after 24 h of incubation, SEM cells cultured in CSFmax significantly increased consumption of glutathione, as well as of cystine, which reached significance by 72 h of incubation (with ∼80% of available metabolite being consumed; Figure 2A, Supplementary Figure 2E-F). These two metabolites are known to play a crucial role in combating oxidative stress [33], hence indicating a response to redox imbalance.

**Figure 2:**
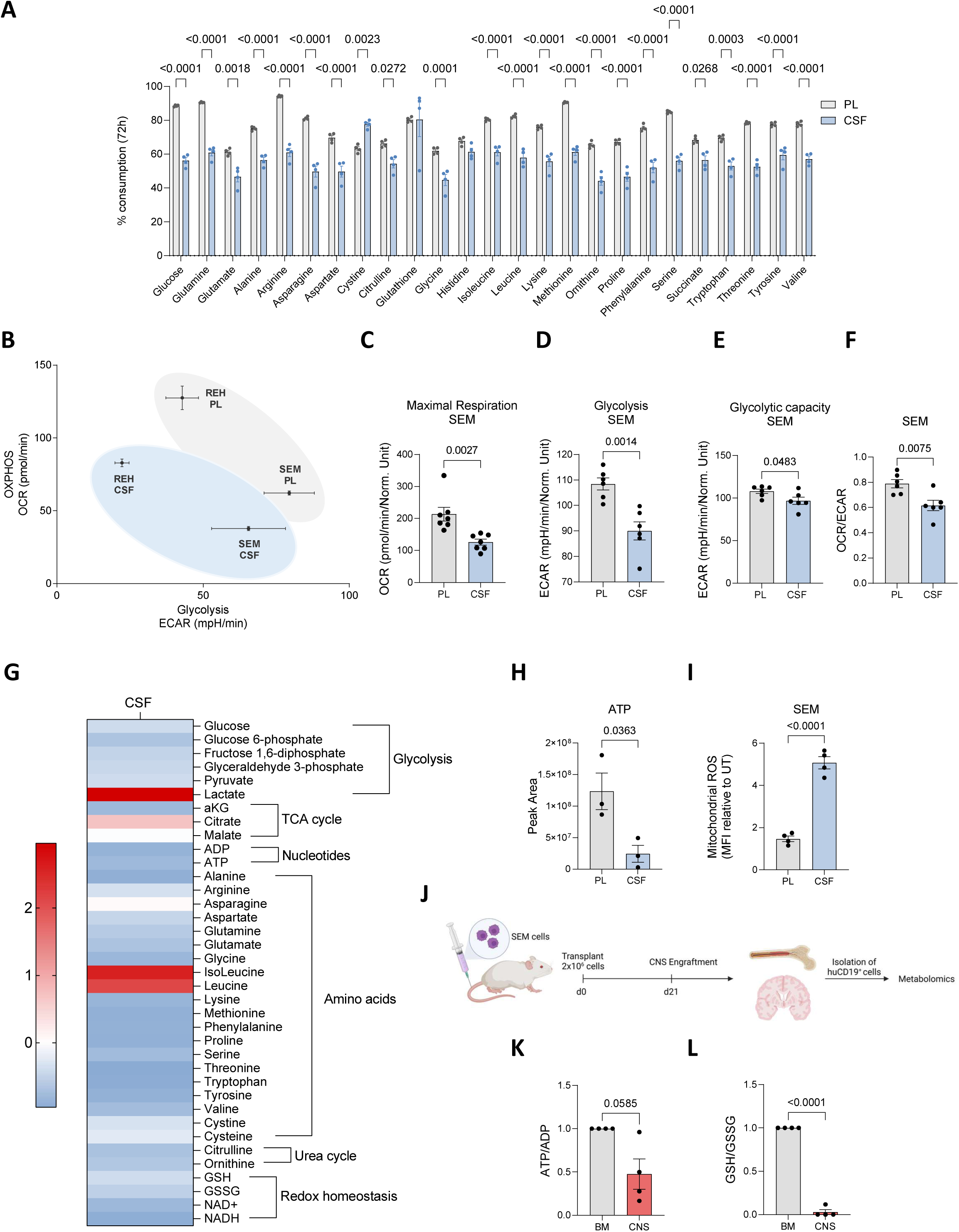
Culture of ALL cells in CSFmax extensively alters their metabolic activity comparably to CNS-derived ALL cells. **A,** Percentage of consumption of metabolites in SEM cells cultured in Plasmax (PL) and CSFmax (CSF) for 72 h (n=4 independent cultures). **B,** Metabolic phenotype map of SEM and REH cells cultured in Plasmax (PL) and CSFmax (CSF) for 24 h (n=6-7 independent wells, representative of two-three independent experiments). **C,** Quantification of maximal respiration in SEM cells in response to culture conditions as indicated in (B) (n=7 independent wells, representative of three independent experiments). **D,E,** Quantification of glycolysis (D) and glycolytic capacity (E) in SEM cells in response to culture conditions as indicated in (B) (n=6 independent wells, representative of two independent experiments). **F,** OCR/ECAR ratio in SEM cells in response to culture conditions as indicated in (B) (n=6-7 independent wells, representative of two-three independent experiments). **G,** Heatmap showing fold change of intracellular levels of glycolysis, TCA cycle, nucleotides, and other relevant metabolites in SEM cells cultured in CSFmax (CSF) compared to SEM cells cultured in Plasmax (PL) for 24 h (n=3 independent cultures). **H,** Total intracellular levels of ATP in SEM cells in response to culture conditions as indicated in (B) (n=3 independent cultures). **I,** Quantification of relative mean fluorescence intensity (MFI) of mitochondrial ROS in SEM cells in response to culture conditions as indicated in (B) (n=4 independent experiments). **J,** Schematic diagram of experimental methodology for *in vivo* targeted metabolomics assay. Four female NSG immunodeficient mice of age 6-10 weeks were transplanted with SEM cells and at experimental endpoint human CD19^+^ cells were isolated from the BM and the brain for metabolomic analysis through LC-MS. **K, L,** Intracellular ATP/ADP (K) and GSH/GSSG (L) ratio relative to SEM cells isolated from BM (n=4 biological replicates). Data are shown as the mean ± s.e.m. P-values were calculated with a repeated measure two-way ANOVA with Sidak’s multiple comparison test (A), unpaired t-test (C, D, E, F, H, I), and paired t-test (K, L).

By measuring O_2_ consumption rate (OCR) and extracellular acidification rate (ECAR), we showed that SEM and REH cells cultured in CSFmax reduced mitochondrial respiration, as previously reported in CNS-derived leukaemic cells [25], and glycolysis compared to cells cultured in Plasmax (Figure 2B-E, Supplementary Figure 3A-G). Notably, SEM cells cultured in CSFmax exhibited a significant reduction in OCR/ECAR ratio (Figure 2F), indicating a preferential utilization of glycolysis. Metabolomic analysis revealed an overall reduction in intracellular levels of most amino acids in SEM cells cultured in CSFmax, compared to Plasmax (Figure 2G), consistent with the reduced consumption of amino acids (Figure 2A). While SEM cells cultured in CSFmax suggested greater dependency on glycolysis (Figure 2F), we observed a downward trend in the intracellular levels of glucose and other glycolytic intermediates, such as glucose-6-phosphate, fructose-1,6-diphosphate, glyceraldehyde-3-phosphate, and pyruvate, in SEM cells cultured in CSFmax (Figure 2G, Supplementary Figure 4A-B). Interestingly, intracellular lactate levels were significantly increased in SEM cells cultured in CSFmax (Figure 2G, Supplementary Figure 4C), further suggesting the uptake of lactate to support their survival, a well-recognised survival mechanism utilised by cells within the human brain and by cancer cells found in hypoxic niches [34, 35].

Moreover, SEM cells cultured in Plasmax and CSFmax did not show any differences in intracellular levels of TCA cycle intermediates, citrate and malate, however a significant reduction in intracellular α-ketoglutarate (AKG) levels was observed in SEM cells cultured in CSFmax, illustrating partial impairment in TCA cycle activity (Figure 2G, Supplementary Figure 4D). Importantly, we showed that culture of SEM cells in CSFmax significantly reduced intracellular levels of ATP (Figure 2H), signifying reduced energy levels. Furthermore, SEM cells had lower intracellular levels of glutathione, both in its reduced (GSH) and oxidized (GSSG) form, although this did not reach statistical significance (Figure 2G, Supplementary Figure 4E-F). The levels of mitochondrial ROS in SEM cells cultured in CSFmax were significantly elevated (Figure 2I), strengthening the concept that cells in CSF undergo sustained oxidative stress.

To examine the metabolic changes ALL cells undergo within the CNS microenvironment, 2×10^6^ SEM cells were transplanted in NSG immunodeficient mice (Figure 2J). Metabolomic analysis on human CD19^+^ cells isolated from the BM and brain revealed reduced energy levels in CNS-derived leukaemic cells when compared to BM-derived leukaemic cells, indicated by a reduction in the ATP/ADP ratio (Figure 2K). Importantly, we observed a significant decrease in the ratio of reduced to oxidised glutathione (GSH/GSSG) intracellular levels in CNS-derived leukaemic cells when compared to BM-derived leukaemic cells (Figure 2L), further suggesting that within the CNS microenvironment ALL cells are under greater oxidative stress.

### Autophagy associated genes are upregulated in CNS-ALL revealing a dependency on mitophagy receptor BNIP3 for CNS engraftment

We next aimed to shed light on how ALL cells maintain long-term survival within the CNS microenvironment. Further examination of the transcriptional adaptations of ALL cells in the CNS microenvironment through over-representation analysis revealed various gene sets to be enriched in the CNS compared to the periphery (spleen), including autophagy of mitochondrion (mitophagy) and response to hypoxia (Figure 3A). Specifically, we identified multiple autophagy and hypoxia associated genes to be significantly upregulated in CNS-derived leukaemic cells (Figure 3B-C). While no significant transcriptional differences were observed in genes involved in the canonical autophagy pathway, such as Unc-51 Like Autophagy Activating Kinase 1 (ULK1) and autophagy related 7 (ATG7), we did observe significant transcriptional upregulation of hypoxia-induced mitophagy receptors, BCL2 Interacting Protein 3 (BNIP3) and BCL2 Interacting Protein 3 Like (BNIP3L), in CNS-derived leukaemic cells (Figure 3D, Supplementary Figure 5A), suggesting the involvement of receptor-mediated mitophagy in hypoxic CNS-ALL cells [36–38]. Importantly, multiple autophagy genes involved in the canonical autophagy pathway were found to be upregulated in CSF-derived leukaemic cells when compared to BM-derived leukaemic cells directly isolated from patients with B-ALL at the time of combined BM and CNS relapse, including ATG7, as well as mitophagy receptor BNIP3 (Figure 3E, Supplementary Figure 5B-C). Therefore, we hypothesised that ALL cells may upregulate and utilise the autophagy machinery to support their survival in the CNS microenvironment.

**Figure 3:**
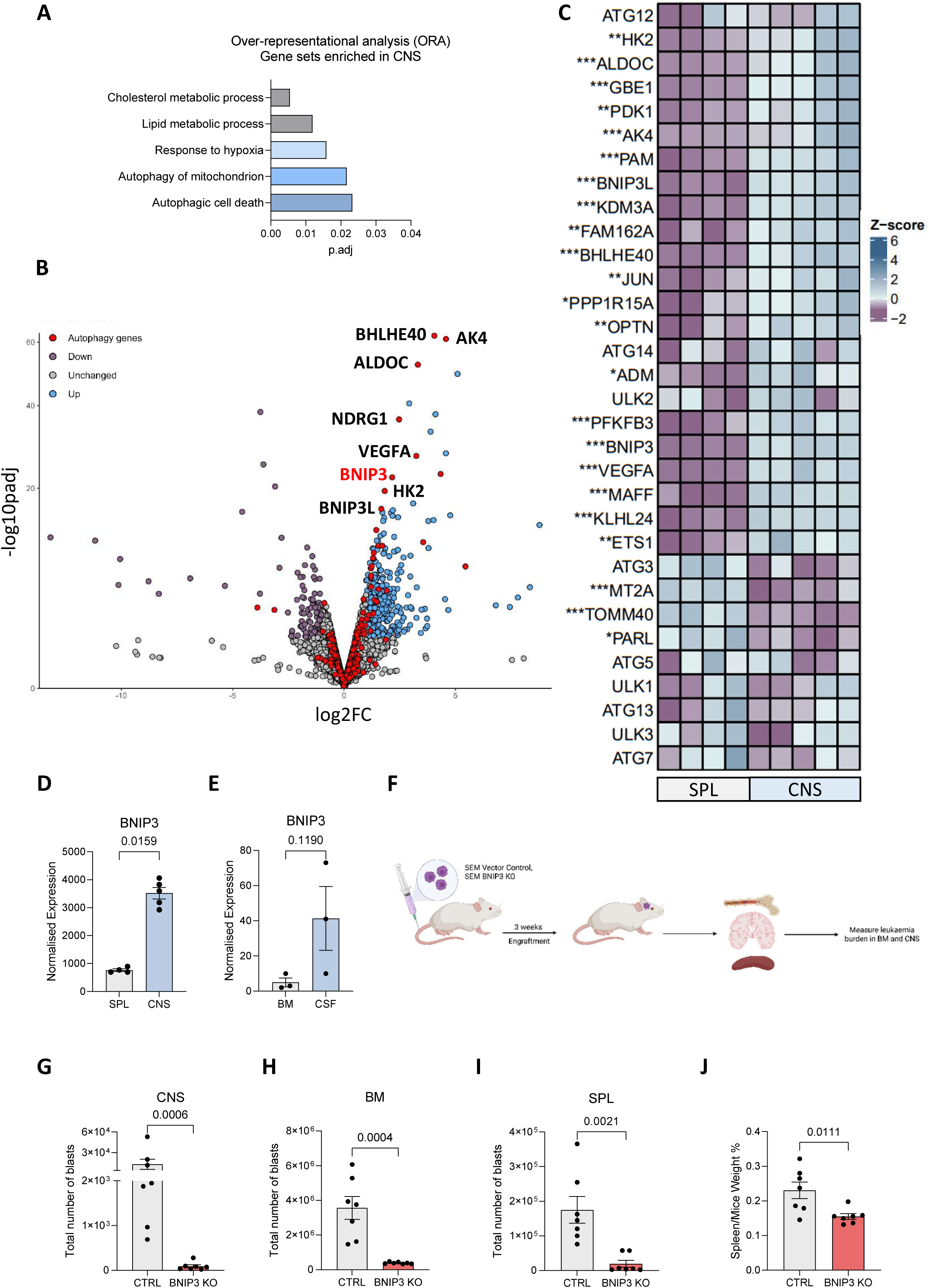
Autophagy associated genes are upregulated in CNS-derived ALL cells and BNIP3 inhibition reduces CNS leukaemia burden. **A,** Over-representational analysis (ORA) of gene sets enriched in human SEM and REH cells isolated from the CNS compared to human SEM and REH cells isolated from the spleen of xenografted mice (n=2 independent groups of 5 mice for SEM cells, n=2-3 independent groups of 5 mice for REH cells). Publicly available dataset used: GSE135115. **B,** Volcano plot representing fold change of transcriptional expression in response to conditions as indicated in (A). Upregulated genes are in blue, downregulated genes in purple, non-significant genes in grey, and autophagy associated genes in red. Plot includes names of autophagy associated genes that are significantly upregulated. Genes were classified as significantly differentially expressed when p<0.05 and log_2_FoldChange ≥ 2. **C,** Heatmap representation of differentially expressed genes between conditions as indicated in (A). P-values: * p>0.05, ** p>0,01, *** p>0.001. Purple indicates upregulation of genes and blue indicates downregulation of genes. **D,** Normalised expression values of BNIP3 in human SEM and REH cells isolated from the CNS and the spleen. Publicly available dataset used: GSE135115. **E,** Normalised expression values of BNIP3 in human ALL cells isolated from the CSF and the BM of patient-derived samples with B-ALL at the time of combined BM and CNs relapse (n=3 biological replicates). Publicly available dataset used: GSE81519. **F,** Schematic diagram of experimental methodology for cell line xenograft experiment. Fourteen female NSG immunodeficient mice were transplanted with either SEM vector control (CTRL) or SEM BNIP3 knockout (KO) cells to measure leukaemia burden in the BM, CNS, and spleen. **G, H, I,** Number of human CD19^+^ SEM CTRL or SEM BNIP3 KO cells harvested from the brain (G), BM (H), and spleen (I) at experimental endpoint (n=7 biological replicates for CTRL group; n=7 biological replicated for BNIP3 KO group). **J,** Spleen/body weight ratio of the mice at experimental endpoint (n=7 biological replicates for CTRL group; n=7 biological replicated for BNIP3 KO group). Data are shown as the mean ± s.e.m. P-values were calculated with unpaired t-test (D, E, G, H, I, J).

We firstly aimed to explore the role of the mitophagy receptor BNIP3 in CNS-ALL. Notably, SEM cells cultured in CSFmax, in both normoxia and hypoxia, revealed a clear increase in HIF-1a, BNIP3, and BNIP3L protein levels in hypoxia (Supplementary Figure 6A), as well as when cultured in RPMI and Plasmax, suggesting that this may not be CNS-specific. To test whether genetic inhibition of BNIP3 would impair CNS engraftment, we generated SEM BNIP3 knockout (KO) cells (Supplementary Figure 6B). Next, control and BNIP3 KO cells were transplanted in NSG immunodeficient mice and BM, brain, and spleen harvested when mice reached the overall severity threshold (Figure 3F). Despite variations in CNS infiltration levels in mice xenografted with control cells, flow cytometry analysis (Supplementary Figure 6C) demonstrated that loss of BNIP3 impaired the engraftment capacity of ALL cells, measured by significant reduction (>95%) in CNS tumour load (Figure 3G), with similar results obtained in BM and spleen (Figure 3H-J).

### CSFmax culture induces autophagy and selectively sensitises ALL cells to autophagy inhibition *in vitro*

Restricted by the lack of BNIP3 inhibitors along with the lack of a CNS-specific phenotype, we decided to broaden our approach and asked whether autophagy may play a role in the adaptation of ALL cells in the CNS microenvironment. Subsequently, we aimed to assess whether inhibiting the canonical autophagy pathway (which may affect both the clearance of mitochondria and of nutrients/amino acids) is equally detrimental for CNS-ALL. Firstly, we generated stable ALL cell lines expressing GFP-LC3-RFP-LC3ΔG (Supplementary Figure 6D) [39], a fluorescent autophagic flux probe. This allows robust assessment of autophagic degradation in SEM and REH cells cultured in CSFmax, with or without bafilomycin treatment, an autophagy inhibitor which acutely blocks autophagic flux by inhibiting the autophagosome and lysosome fusion [40] (Figure 4A). Reassuringly, we observed that both SEM and REH cells expressing GFP-LC3-RFP-LC3ΔG cultured in CSFmax exhibited significantly higher autophagy flux compared to when cultured in RPMI or Plasmax, indicated by the significant reduction in the GFP/RFP ratio (Figure 4B-C). Importantly, bafilomycin reversed this effect and blocked autophagy flux, observed by the increased GFP/RFP ratio (Figure 4B-C). Following this, we measured the protein levels of p62 and LC3B, two key autophagy markers that get degraded during autophagy (Figure 4A). Indeed, we observed autophagy induction in SEM cells cultured in CSFmax, indicated by the reduction in p62 protein levels and modest increase in lipidated LC3-II levels (Supplementary Figure 6A). Bafilomycin treatment in SEM cells either cultured in CSFmax in normoxia or hypoxia led in the accumulation of both p62 and LC3-II levels when compared to untreated cells (Supplementary Figure 6A), excluding accumulation of autophagosomes due to defective autophagosome-lysosome fusion [41, 42], and confirming high autophagy flux in cells cultured in CSFmax. Notably, we observed a similar reduction in p62 levels in SEM cells cultured in CSFmax either in normoxia or hypoxia, when compared to RPMI and Plasmax-cultured cells (Supplementary Figure 6A), highlighting that the increased autophagy flux in CSFmax-cultured cells is mainly driven by reduction in nutrients and occurs independently of O_2_ levels. Similarly to CSFmax-cultured cells, we validated that autophagy is induced in CNS-derived leukaemic cells isolated from two NSG mice xenografted with SEM vector control cells, observed by the reduction in p62 protein and LC3-I levels when compared to BM-derived leukaemic cells (Figure 4D), along with higher accumulation of lipidated LC3-II when compared to LC3-I levels in CNS-derived leukaemic cells (Figure 4D).

**Figure 4:**
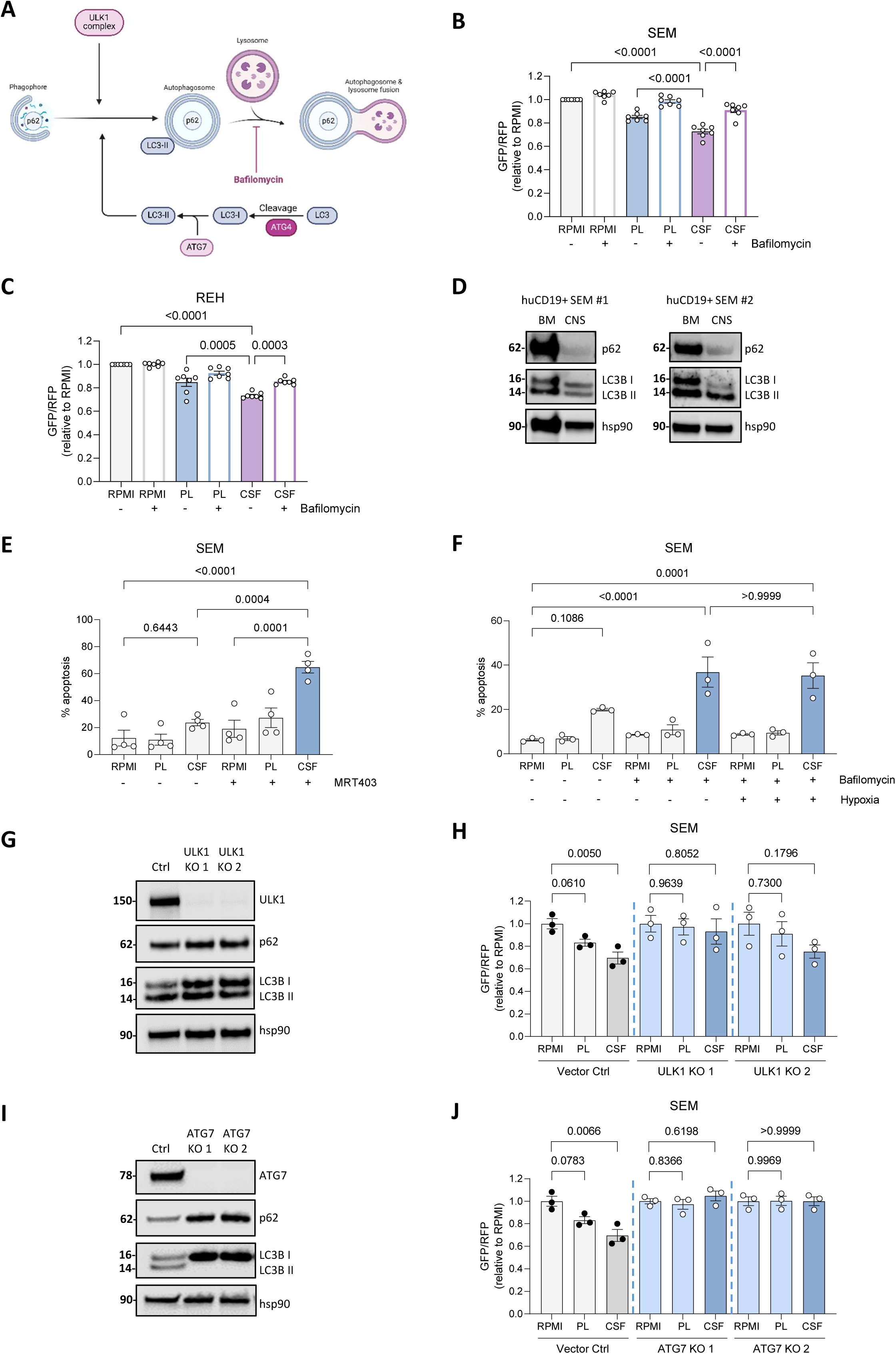
Autophagy flux is higher in ALL cells cultured in CSFmax. **A,** Schematic diagram of autophagy pathway and the mechanism of action of bafilomycin A1 to inhibit autophagy. **B, C,** Relative GFP/RFP ratio of mean fluorescence intensity (MFI) in SEM (C) and REH (D) cells expressing GFP-LC3-RFP-LC3ΔG autophagic flux probe cultured in RPMI, Plasmax (PL), and CSFmax (CSF) for 24 h with or without 50 nM of bafilomycin A1 treatment (n=7 independent experiments). **D,** Immunoblot analysis showing changes in autophagy markers in human CD19^+^ (huCD19^+^) SEM cells isolated from the bone marrow (BM) and the brain (CNS) (n=2 biological replicates). **E,** Percentage of apoptosis in SEM cells cultured in RPMI, Plasmax (PL), and CSFmax (CSF) for 24 h in normoxia (21% O_2_) with or without 1 μM of MRT403 treatment (n=3 independent experiments). **F,** Percentage of apoptosis in SEM cells cultured in RPMI, Plasmax (PL), and CSFmax (CSF) for 24 h in normoxia (21% O_2_) or hypoxia (1% O_2_) with or without 50 nM of bafilomycin A1 treatment (n=3 independent experiments). **G,** Immunoblot analysis showing deletion of ULK1 and changes in autophagy markers in SEM cells cultured in RPMI. **H,** Relative GFP/RFP ratio of mean fluorescence intensity (MFI) in SEM vector control (CTRL) and ULK1 knockout (KO) single clone 1 and 2 cells expressing GFP-LC3-RFP-LC3ΔG autophagic flux probe cultured in RPMI, Plasmax (PL), and CSFmax (CSF) for 24 h (n=3 independent experiments). **I,** Immunoblot analysis showing deletion of ATG7 and changes in autophagy markers in SEM cells cultured in RPMI. **J,** Relative GFP/RFP ratio of mean fluorescence intensity (MFI) in SEM CTRL and ATG7 KO single clone 1 and 2 cells expressing GFP-LC3-RFP-LC3ΔG autophagic flux probe cultured in RPMI, Plasmax (PL), and CSFmax (CSF) for 24 h (n=3 independent experiments). Data are shown as the mean ± s.e.m. P-values were calculated with a repeated measure one-way ANOVA with Tukey’s multiple comparison test (B, C, E, F, H, J).

Next, we tested the sensitivity of ALL cells upon pharmacological inhibition of autophagy by inhibiting the initiation step of the pathway with the ULK1 inhibitor MRT403 [43] (Supplementary Figure 6E), and bafilomycin. Indeed, pharmacological inhibition of autophagy treatment significantly increased the percentage of apoptosis of SEM and REH cells cultured in CSFmax (Figure 4E-F, Supplementary Figure 6F-G). Importantly, neither inhibitor sensitised SEM and REH cells cultured in RPMI or Plasmax, highlighting that ALL cells rely on autophagy selectively when cultured in CSFmax. Notably, there was no significant difference in the percentage of apoptosis between ALL cells cultured in CSFmax in normoxia and hypoxia treated with bafilomycin (Figure 4F, Supplementary Figure 6G), further indicating that ALL cells mainly use autophagy for their survival in response to nutrient stress, rather than in response to low O_2_ levels.

Furthermore, we examined the impact of specific inhibition of autophagy at the initiation step and at a later stage of the pathway, using CRISPR-Cas9 technology to generate SEM ULK1 KO and ATG7 KO cell lines (Figure 4G, I). As expected, p62 protein levels were increased together with accumulation of LC3-I and LC3-II (indicating stalled early autophagosomes as previously reported [43, 44], demonstrating autophagy inhibition following ULK1 KO (Figure 4G). As before, vector control cells cultured in CSFmax exhibited higher autophagy flux compared to RPMI or Plasmax cultured cells, indicated by the significant reduction in GFP/RFP ratio (Figure 4H, J). Notably, loss of ULK1 prevented acute increase in autophagy flux in CSFmax, observed by the absence of reduction in the GFP/RFP ratio in ULK1 KO cells cultured in CSFmax, compared to control cells (Figure 4H). Additionally, loss of ATG7 which inhibits the lipidation of LC3-II and hence the conjugation of LC3-II to the autophagosome membrane, showed a clear block of basal autophagy, demonstrated by an increase in p62 protein level and absence of formation of LC3-II (Figure 4I). ATG7 KO also blocked acute autophagy induction following culture in CSFmax assessed by the unaffected GFP/RFP ratio in ATG7 deficient cells (Figure 4J).

### ULK1 and ATG7 ablation decrease engraftment capacity of ALL cells in the CNS

To test whether genetic inhibition of autophagy would impair ALL survival in the CNS microenvironment, three separate cell line xenotransplantation experiments were employed by transplanting NSG immunodeficient mice with SEM vector control, ULK1 KO, and ATG7 KO cells (Figure 5A). Notably, we included both male and female mice to identify potential sex-selective effects. Crucially, loss of both ULK1 and ATG7 impaired the engraftment capacity of SEM cells in the CNS (Figure 5B). Surprisingly, ULK1 and ATG7 deficient cells did not exhibit a significant reduction in BM tumour load in NSG mice when compared to mice xenografted with control cells, although there was a noticeable trend of reduction in BM tumour burden in mice xenografted with ULK1 deficient cells, which reached statistical significance in one of the experiments (experiment #3) (Figure 5C). Interestingly, loss of ULK1 and ATG7 impaired the xenograft capacity of ALL cells in the spleen and significantly reduced the spleen to body weight ratio of the mice at experimental endpoint (Figure 5D-E), in line with results obtained following BNIP3 KO (Figure 3H).

**Figure 5:**
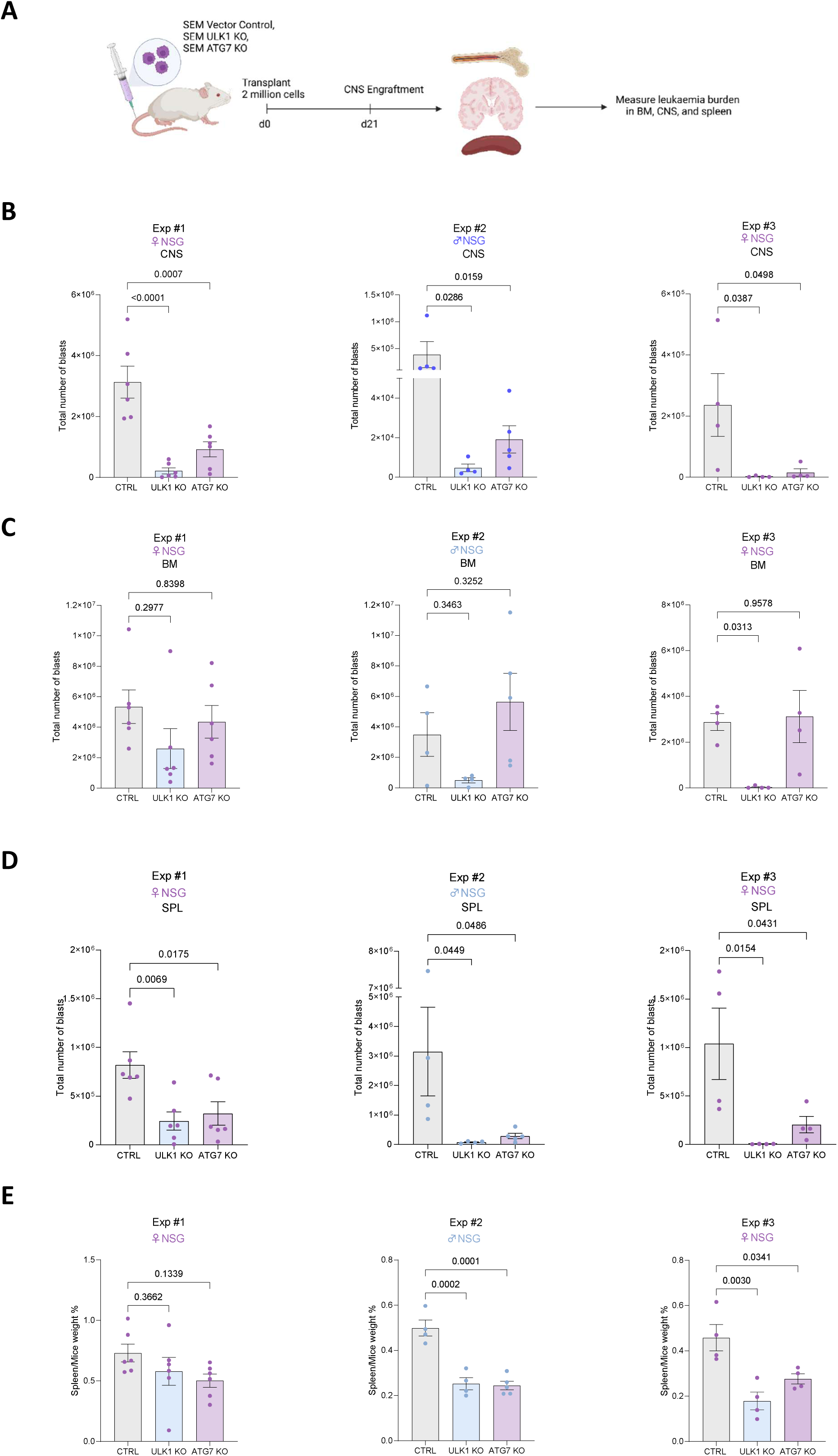
ULK1 and ATG7 deficient cells impair xenograft capacity of ALL cells in the CNS. **A,** Schematic diagram of experimental methodology for cell line xenograft experiments. NSG immunodeficient mice were transplanted with either SEM vector control (CTRL), SEM ULK1 knockout (KO), or SEM ATG7 KO cells to measure leukaemia burden in the BM, CNS, and spleen. **B, C, D,** Number of human CD19^+^ SEM CTRL, SEM ULK1 KO, or SEM ATG7 KO cells harvested from the brain (B), BM (C), and spleen (D) at experimental endpoint (Exp #1: n= 6 mice for CTRL group; n=6 mice for ULK1 KO group; n=6 mice for ATG7 KO group) (Exp #2: n= 4 mice for control group; n=4 mice for ULK1 KO group; n=5 mice for ATG7 KO group) (Exp #3: n= 4 mice for control group; n=4 mice for ULK1 KO group; n=4 mice for ATG7 KO group) **E,** Spleen/body weight ratio of the mice at experimental endpoint (Exp #1: n= 6 mice for CTRL group; n=6 mice for ULK1 KO group; n=6 mice for ATG7 KO group) (Exp #2: n= 4 mice for control group; n=4 mice for ULK1 KO group; n=5 mice for ATG7 KO group) (Exp #3: n= 4 mice for control group; n=4 mice for ULK1 KO group; n=4 mice for ATG7 KO group). Data are shown as the mean ± s.e.m. P-values were calculated with a repeated measure one-way ANOVA with Dunnet’s multiple comparison test (B, C, D, E).

Intrigued by the fact that we did not observe any effect in the BM tumour load upon loss of ATG7, we evaluated the protein levels of ATG7 and of autophagy markers in isolated human CD19^+^ cells retrieved from the BM. As expected, human CD19^+^ cells isolated from the BM of mice xenografted with ATG7 deficient cells had low ATG7 protein levels, increased p62 and LC3-I protein levels, and lack of formation of LC3-II, when compared to human CD19^+^ cells isolated from the BM of mice xenografted with control cells (Supplementary 7A). This confirmed that SEM ATG7 KO cells transplanted in NSG mice were indeed autophagy deficient and prompted us to question how persistent ATG7 deficient cells, unable to support energy and biomass generation through autophagy, may potentially further re-program and survive in the CNS. Therefore, metabolomic analysis was carried out in CNS-derived and BM-derived SEM vector control and ATG7 deficient cells isolated from xenografted mice (metabolomic analysis could not be carried out in ULK1 deficient cells due to low cell numbers). We observed a reduction in the intracellular levels of most amino acids in CNS-derived ATG7 deficient cells when compared to CNS-derived control cells (Supplementary Figure 7B). Interestingly, the intracellular levels of most amino acids remained unchanged in BM-derived ATG7 deficient cells when compared to BM-derived control cells (Supplementary Figure 7B), suggesting that autophagy is important for the maintenance of amino acid levels in ALL cells specifically within the CNS microenvironment, where extracellular levels of amino acids and other nutrients are limited and cells may need to rely more on recycling of nutrients.

Furthermore, intracellular levels of glycolysis related metabolites, such as glucose, glyceraldehyde-3-phosphate, and lactate were increased in CNS-derived control cells, and this increase was further enhanced in CNS-derived ATG7 deficient cells (Supplementary Figure 7B-E). Of note, in the BM, loss of ATG7 did not cause a similar increase in glycolysis related metabolites and lactate as in the CNS (Supplementary Figure 7B-E). These findings either suggest the potential uptake of glucose and increased reliance on glycolysis of ALL cells upon autophagy deficiency specifically within the CNS, or the reduction of flow through the glycolysis pathway leading to the accumulation of glucose and key glycolysis intermediates. Importantly, the overall energy status of CNS-derived leukaemic cells was further decreased upon loss of ATG7, indicated by the enhanced reduction in the intracellular ATP/ADP ratio in CNS-derived ATG7 deficient cells when compared to BM-derived control cells (Supplementary Figure 7F). Although loss of ATG7 in CNS-derived cells exhibited a non-significant decrease in the ATP/ADP ratio, this decreased trend further suggested that ALL cells affect their energy status in the CNS through autophagy. Surprisingly, we observed that the levels of intracellular GSH were increased, specifically in CNS-derived ATG7 deficient cells (Supplementary Figure 7B), possibly reflecting a contribution to compensatory antioxidant response [33].

## Discussion

It is widely recognised that the disproportionate and nonphysiological composition of conventional cell culture media rewires the metabolism of cancer cells and generates discrepancies between *in vitro* and *in vivo* findings [1–4]. The meningeal niche is a prime example of a defined metabolic microenvironment in which the limited nutrient and O_2_ availability in CSF [7, 11–13] is not reflected in any of the high-nutrient composition of commercially available cell culture media. As accumulating evidence shows that the low O_2_ and nutrient availability in the CSF induce metabolic reprogramming in ALL cells [25–27], investigating and targeting metabolic dependencies of ALL blasts in the CNS is an attractive approach to directly eradicate them. However, so far, studies have been primarily supported through animal work due to the lack of a physiological relevant cell culture medium. This hinders the extensive *in vitro* investigation on CNS-disease mechanisms required to improve risk stratification and to reduce doses of neurotoxic intrathecal therapies in childhood CNS-ALL [22, 23]. Here, we developed and systematically formulated a novel physiologic CSF-like cell culture medium (CSFmax), that reflects the nutrient composition of key metabolites detected in human CSF [12].

We demonstrate that culturing ALL cells in CSFmax leads to reduced cell proliferation, slower cell cycle, and decreased OXPHOS activity, a comparable antiproliferative behaviour previously characterised in CNS-derived leukaemic cells [25]. Importantly, we demonstrate that the metabolic activity of CSFmax-cultured ALL cells is similar to CNS-derived leukaemic cells, primarily indicated by the significant reduction in nutrient consumption and reduced intracellular ATP levels, when compared to ALL cells cultured in conventional cell culture media. Additionally, we highlight that CSFmax-cultured ALL cells are under greater oxidative stress, observed by the decreased intracellular GSH levels and accumulation of mitochondrial ROS, both commonly associated with increased oxidative stress [33, 45]. Of note, similarly to human CSF, CSFmax has reduced levels of amino acids required for oxidative stress response, such as cysteine, glycine, and glutamate, the precursors of GSH, highlighting the importance of physiological culture medium in identifying critical antioxidant or stress response mechanisms.

Both autophagy and mitophagy have been previously highlighted as potential therapeutic targets in ALL with evidence suggesting that increased autophagy is linked to chemotherapy resistance in ALL [46–48], however, there is no reported evidence that autophagy plays a role in CNS-ALL. Under CNS hypoxia, HIF-1α activation upregulates hypoxia-responsive genes [25, 49], suggesting BNIP3-mediated mitophagy is a hypoxic stress response [36–38]. Consistent with this, we observed increased BNIP3 and HIF-1α protein levels in ALL cells cultured in CSFmax under hypoxia, indicating hypoxia-induced mitophagy. Indeed, genetic inhibition of BNIP3 drastically reduced the engraftment capacity of ALL cells in the CNS, BM, and spleen, increasing the biological relevance of our *in vitro* findings and indicating that BNIP3-mediated mitophagy is critical for ALL survival *in vivo*.

One caveat is that the impact of BNIP3 KO on CNS disease burden could be due to poor BM engraftment resulting in reduced trafficking to the CNS, or due to a direct impact on survival within the CNS niche. Differentiating these is compounded by a lack of direct or specific BNIP3 inhibitors which could be used once robust engraftment was established. Therefore, we turned our attention to the general autophagy pathway. Notably, there is evidence that targeting autophagy through ULK1 inhibition leads to inhibition of BNIP3-mediated mitophagy [50], also supported by the efficacy of autophagy inhibitors used to modulate BNIP3 activity. Critically, examining the role of the canonical autophagy pathway in CNS-ALL, we demonstrate that ALL cells cultured in CSFmax exhibit increased autophagic degradation and are selectively sensitised to pharmacological inhibition of autophagy, when compared to ALL cells cultured in conventional cell culture media. Considering that the BM acts as a protective niche for ALL cells and contributes to leukaemic cell proliferation, survival, and resistance to therapy [51], our findings suggest that ALL cells residing in the spleen are under greater stress, similarly to ALL cells residing in the CNS, indicating the increased reliance of ALL cells on autophagy under comparable stressful conditions. While this is likely to reflect the importance of recycling nutrients, this dependency may, at least in part, be mediated by mitophagy.

Furthermore, we suggest that the low levels of extracellular amino acids are detrimental to ALL cells upon autophagy inhibition, specifically in the CNS. Subsequently, we demonstrate increased intracellular levels of glycolysis related metabolites, including glucose and lactate in CNS-derived leukaemic cells, which is further enhanced in CNS-derived ATG7 KO cells, whilst this effect is not observed in the BM upon loss of ATG7. These findings, along with the marked reduction in intracellular ATP/ADP ratio in CNS-derived ATG7 deficient cells may reflect a lack of efficient mitophagy, leading to accumulation of dysfunctional mitochondria. Indeed, mitophagy induced by starvation or hypoxia has been shown to be suppressed by loss of ATG7 [52], allowing cells to further reprogram through increased reliance on glycolysis. However, while this encourages design of autophagy inhibitors that can effectively cross the blood–brain barrier, the fact that autophagy may be implicated in neuronal health should be taken into consideration in future studies to avoid increased risk of toxicity in the brain.

Overall, we demonstrate that utilising CSFmax provides deeper insight into how ALL cells interact with the unique environment of the CNS, enabling the identification of previously unrecognised metabolic rewiring mechanisms that support their adaptation and survival within this niche. This may prove critical for the development of future novel therapeutic strategies for CNS leukaemia, other leptomeningeal-tropic malignancies, and CNS autoimmune diseases. Finally, CSFmax may also serve as a valuable tool for advancing our understanding of immune function within the CNS niche, particularly as increasing attention is directed towards the use of CAR-T and CAR-iNKT therapies [53, 54].

## Acknowledgements

The authors would like to thank the Core Services and Advanced Technologies at the Cancer Research UK Scotland Institute (C596/A17196; A31287). We thank Karen Dunn and the Cancer Research UK Scotland Institute mouse facility staff for housing of mice and help with xenograft experiments. We thank Tom Gilbey for cell sorting. We thank ERASCA for providing MRT403 (ULK1 inhibitor). Schematic graphs in Figures 2J, 3F, 4A, 5A and Supplementary Figures 6D, 6E were created using BioRender (https://biorender.com). This work was supported by Himonas PhD Studentship (to E.H.), Cancer Research UK (A29754 to G.V.H.), Blood Cancer UK (18006 to G.V.H.), The Howat Foundation (to G.V.H.), Friends of Paul O’Gorman Leukaemia Research Centre (to G.V.H.), CCLG Little Princess Trust Project Grant CCLGA (2020-2024 to C.H. and A.M.), Cancer Research UK (DRCPFA-Nov21\100001 to C.H.), Cancer Research UK (A23982 to S.T.), and Cancer Research UK core funding of the CRUK Scotland Institute (to D.S.^2^).

## Author contributions

E.H. and G.V.H. conceived and designed the study. E.H. performed all *in vitro* and *in vivo* experiments, analysed and interpreted all data. G.V.H. and C.H. supervised the entire study. A.M., K.R.^1^, M.M.Z. assisted with *in vivo* experiments. K.R.^1^ assisted with bioinformatic analysis. D.S.^1^ and A.I. assisted in generating stable cell lines. D.S.^2^ performed the mass spectrometry. K.R.^2^ assisted in cell culture medium preparation and analysis of extracellular metabolomics experiment. E.H., S.T., C.H., G.V.H. conceived the CSFmax formulation. S.T. provided supervision and material support. E.H. and G.V.H. wrote the manuscript and all other authors reviewed it.

