## Supplementary Figures and Tables for "Physiological cerebrospinal fluid like medium reveals autophagy dependency of leukaemia in the central nervous system"

### Supplementary Figure 1

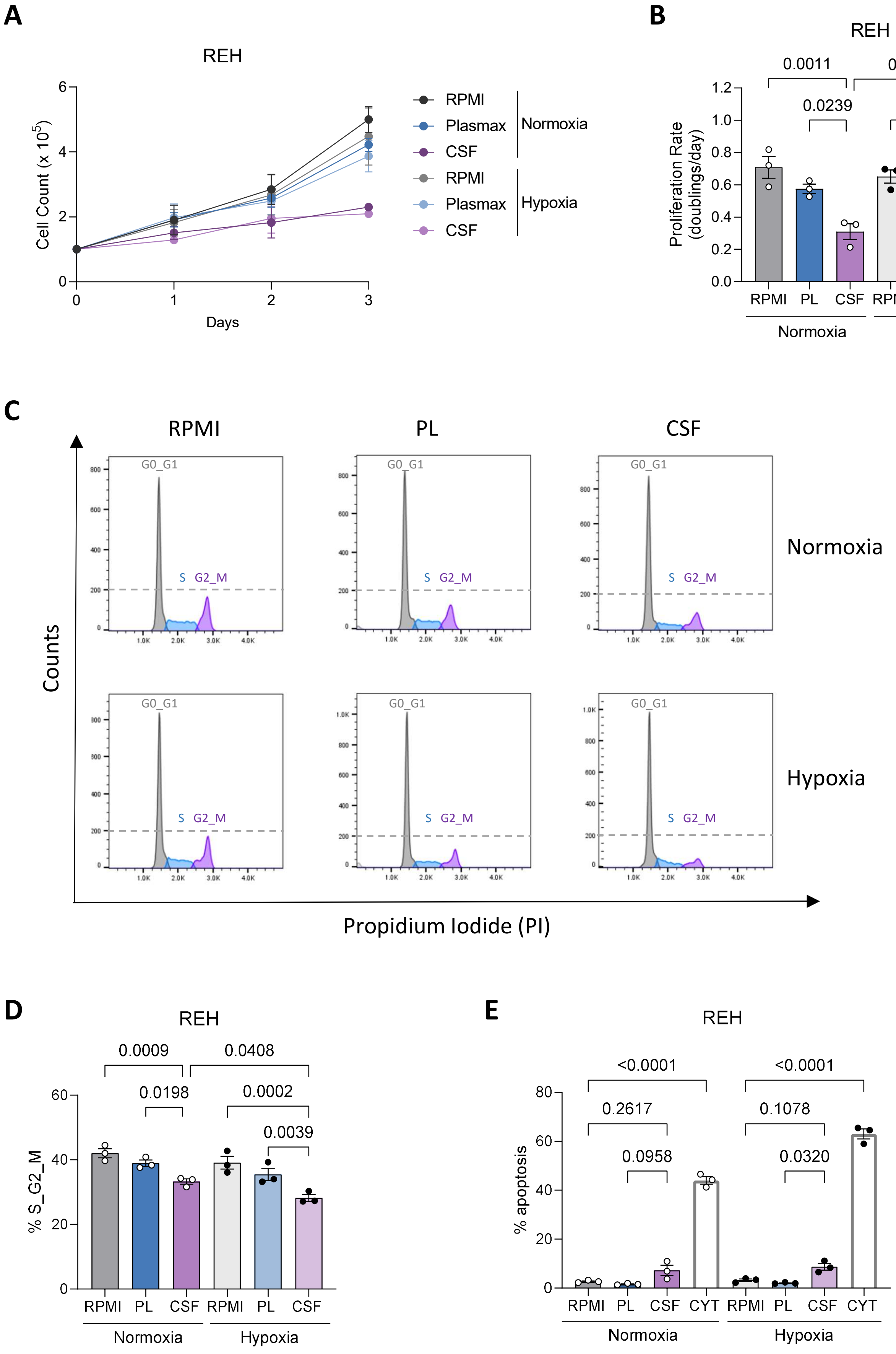

#### Supplementary Figure 2

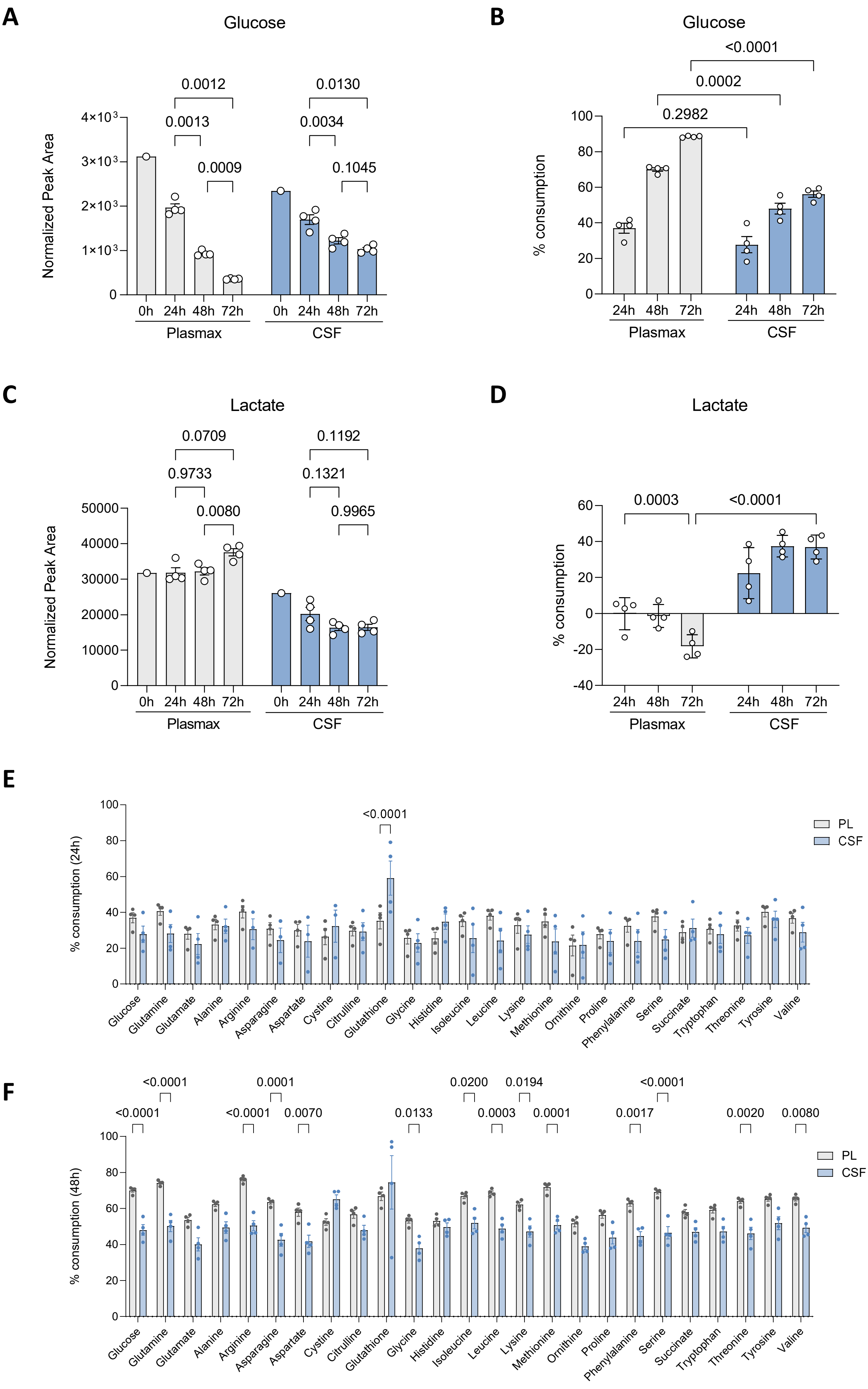

### Supplementary Figure 3

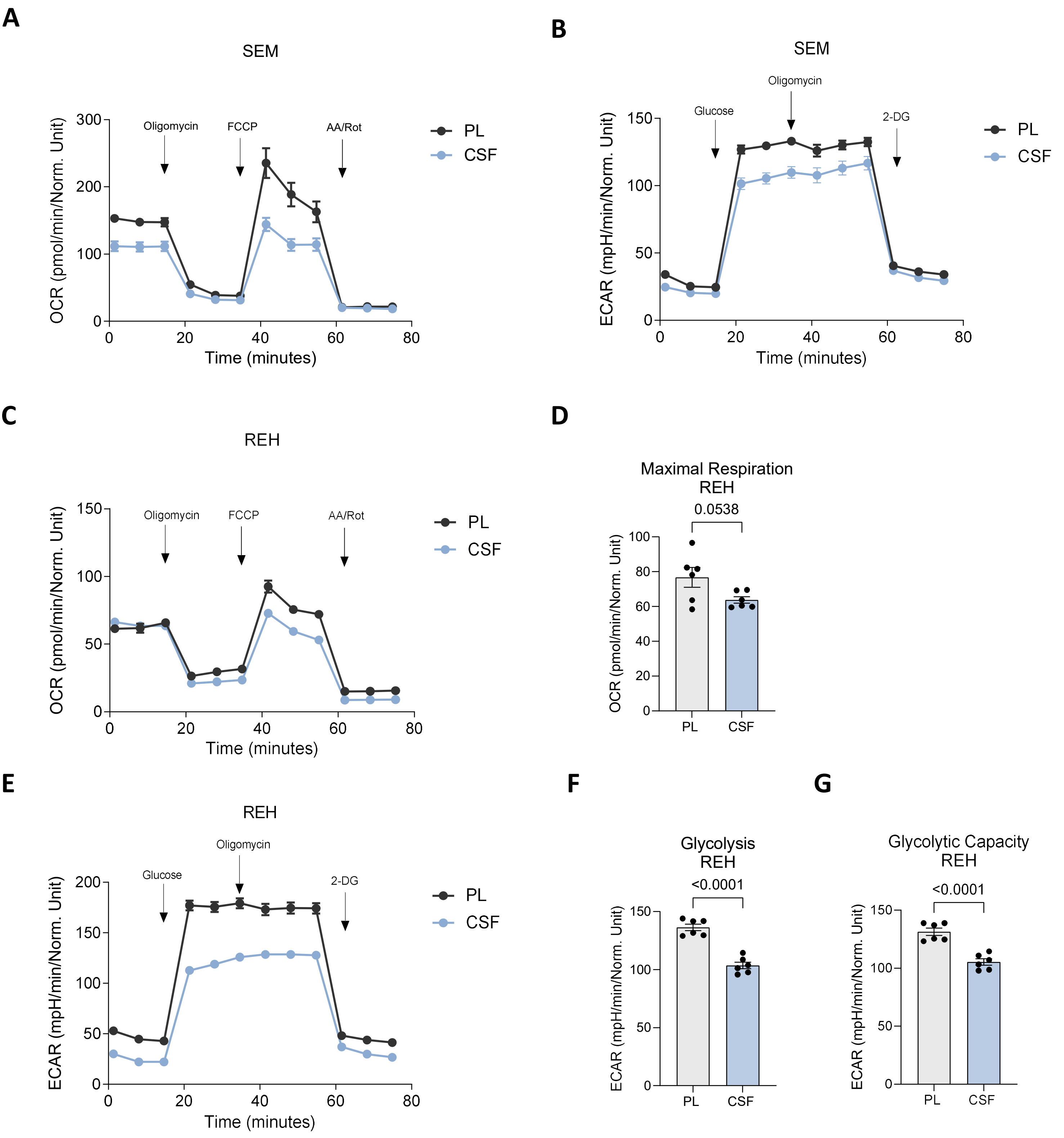

### Supplementary Figure 4

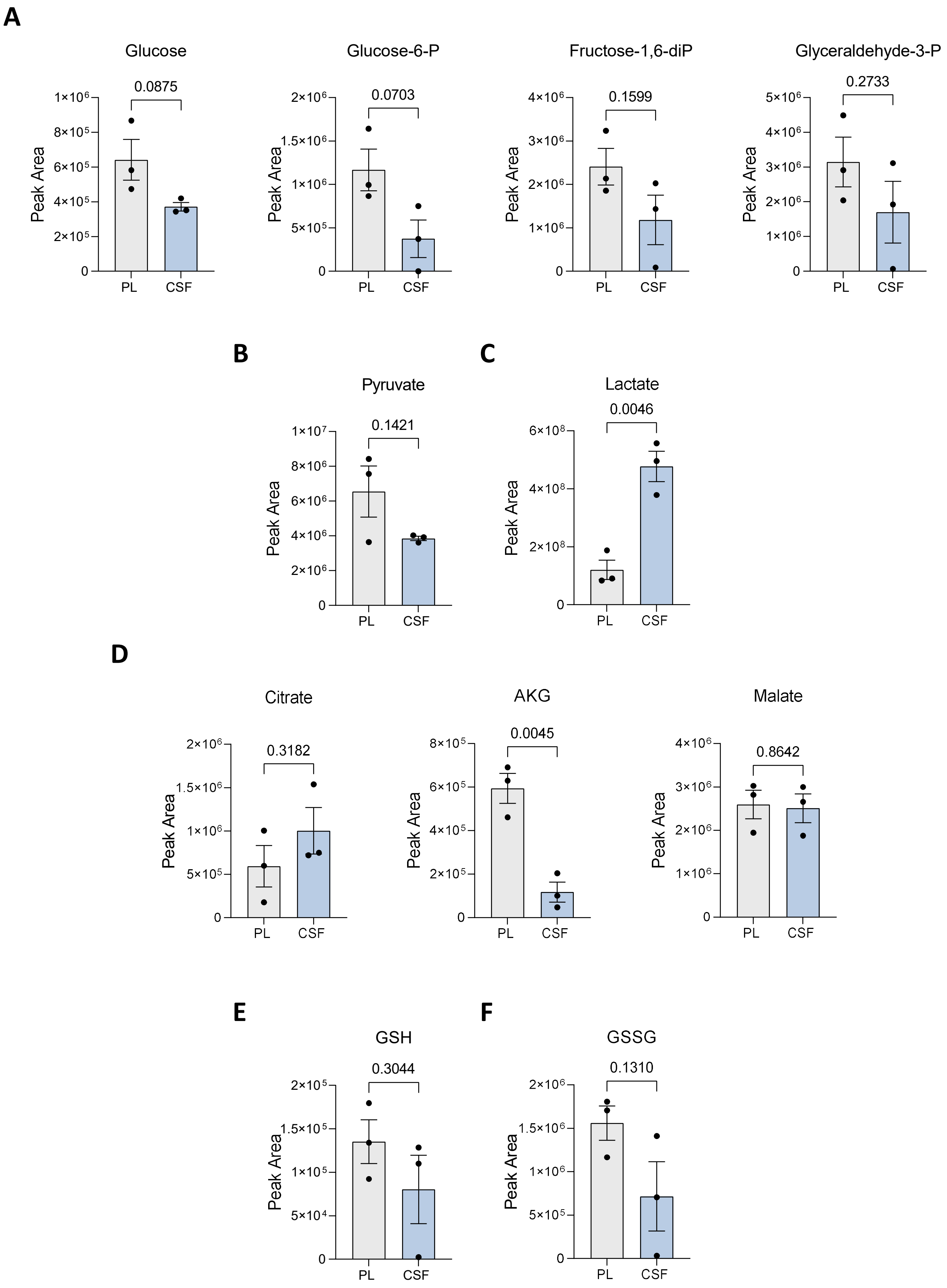

**Supplementary Figure 5**

**A**

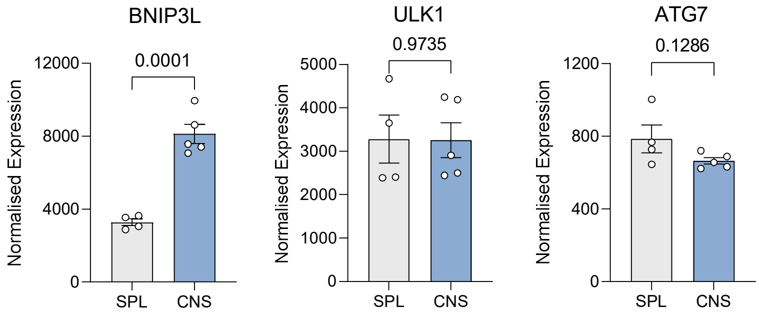


**B**

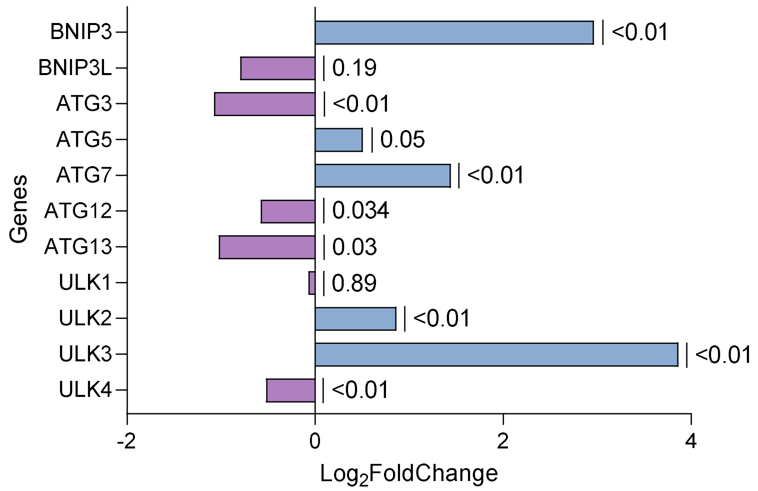


**C**

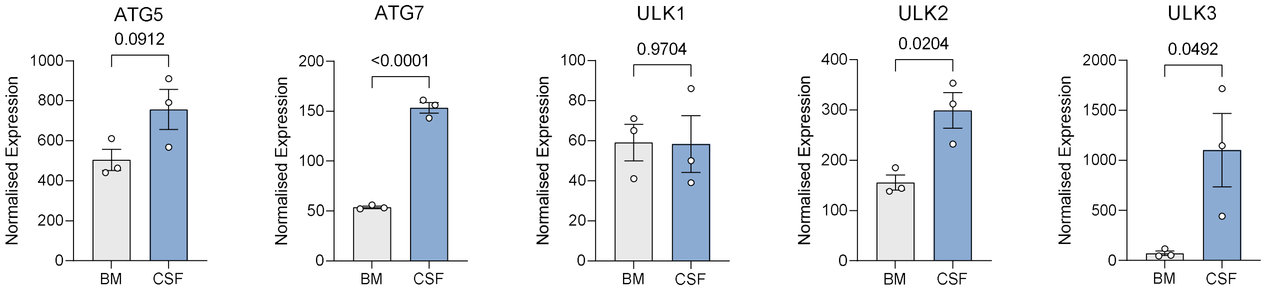

### Supplementary Figure 6

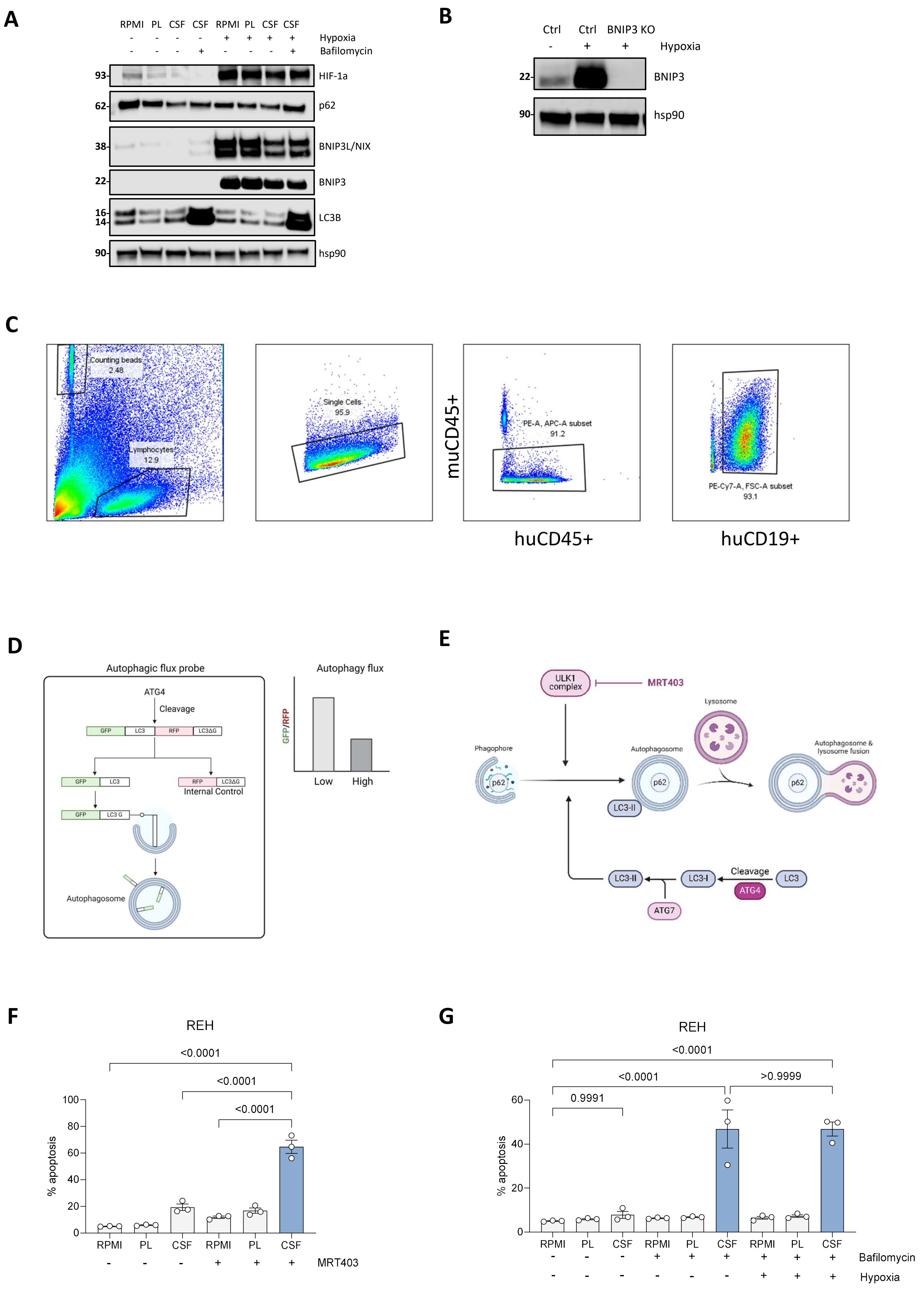

### Supplementary Figure 7

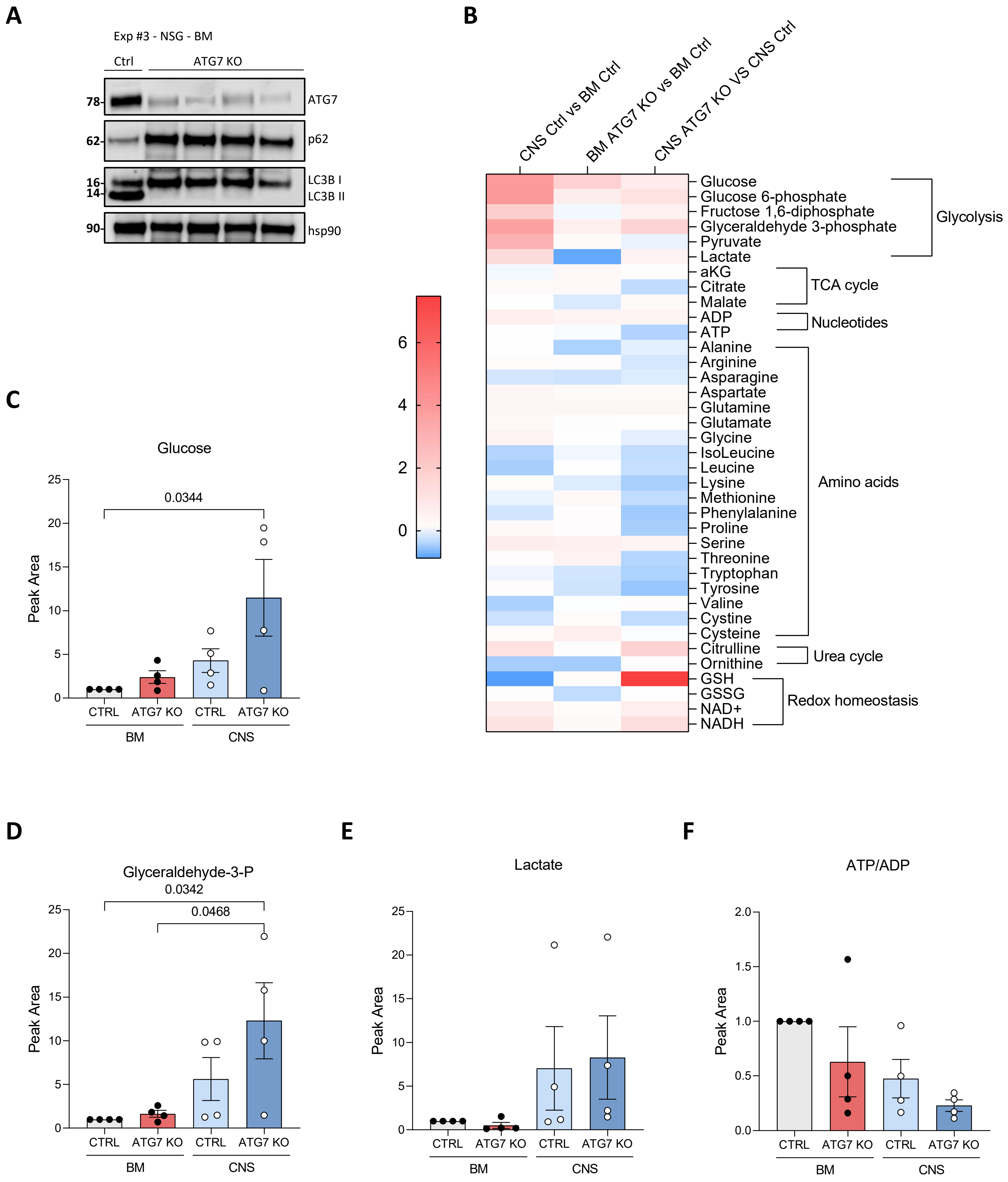

### Supplementary Table 1

| Components: | Concentration (µM): | | | |
|---|---|---|---|---|
| | RPMI | Plasmax | Average CSF metabolome | CSFmax |
| L-Alanine | 0 | 510 | 27,5 | 51 |
| L-Arginine | 1149 | 64 | 19,3 | 19,3 |
| L-Asparagine | 378 | 41 | 6,6 | 6,6 |
| L-Aspartic acid | 150 | 6 | 1,5 | 1,5 |
| L-Glutamate | 136 | 98 | 8,64 | 9,8 |
| Glycine | 133 | 330 | 7,6 | 33 |
| L-Histidine | 97 | 120 | 15,8 | 15,8 |
| L-Isoleucine | 382 | 140 | 7,1 | 14 |
| L-Leucine | 382 | 170 | 13,6 | 17 |
| L-Lysine | 219 | 220 | 26,1 | 26,1 |
| L-Methionine | 101 | 30 | 4 | 4 |
| L-Phenylalanine | 91 | 68 | 11,3 | 11,3 |
| L-Proline | 174 | 360 | 3 | 36 |
| L-Serine | 286 | 140 | 30 | 30 |
| L-Threonine | 168 | 240 | 31 | 31 |
| L-Tryptophan | 25 | 78 | 6 | 7,8 |
| L-Tyrosine | 111 | 74 | 11,3 | 11,3 |
| L-Valine | 171 | 230 | 19,2 | 23 |
| L-Citrulline | N/A | 55 | 3,1 | 5,5 |
| L-Cystine | 208 | 65 | 0,2 | 6,5 |
| L-Ornitine | N/A | 80 | 5,8 | 8 |
| Taurine | N/A | 130 | 8 | 13 |
| Lactate | N/A | 500 | 1608 | 50 |
| D-Glucose | 11111 | 5560 | 3357 | 3357 |
| L-Glutamine | 2055 | 650 | 500 | 500 |

#### Supplementary Table 2

| Supplements: | RPMI | Plasmax | CSFmax |
|---|---|---|---|
| **Serum** | 10% FBS | 10% Dialysed FBS | 5% Dialysed FBS |
| **Lipids** | + | + | - |

**Supplementary Table 3**

| Target | Guide sequence |
|---|---|
| BNIP3 | 5′-CATCACCTACCCCAATCCGA-3′ |
| ULK1 | 5′-AGCAGATCGCGGGCGCCATG-3′ |
| ATG7 | 5′-GAAGCTGAACGAGTATCGGC-3′ |

#### Supplementary Table 4

| Product | Manufacturer | Catalogue Number |
|---|---|---|
| | **Primary antibodies** | |
| ATG7 | Cell Signaling Technology | 8558 (1:1000) |
| BNIP3 | Cell Signaling Technology | 44060 (1:1000) |
| BNIP3L/NIX | Cell Signaling Technology | 12396 (1:1000) |
| HIF-1α | Proteintech | 20960-1 (1:1000) |
| HSP90 | Proteintech | 60318-1 (1:5000) |
| LC3B | Cell Signaling Technology | 2775 (1:1000) |
| P62 | BD Biosciences | 610833 (1:1000) |
| ULK1 | Cell Signaling Technology | 8054 (1:1000) |
| | **Secondary antibodies** | |
| Anti-mouse IgG, HRP-linked | Cell Signaling Technology | 7076 (1:5000) |
| Anti-rabbit IgG, HRP-linked | Cell Signaling Technology | 7074 (1:5000) |
