## Supplemental Materials and Methods_Figure and Table legends for "Physiological cerebrospinal fluid like medium reveals autophagy dependency of leukaemia in the central nervous system"

**Supplementary Figures**

**Supplementary Figure 1: CSFmax culture recapitulates the physiology of ALL cells in the CNS microenvironment. A,B,** Growth curves (A) and proliferation rate (B) of REH cells cultured in RPMI, Plasmax (PL), and CSFmax (CSF) in normoxia (N, 21% $O_2$) or hypoxia (H, 1% $O_2$) (n=3 independent experiments). **C,** Representative flow cytometry plots of SEM cells showing staining with propidium iodide (PI) to assess the effect of cells on cell cycle in response to culture conditions as indicated in (A, B) for 72 h. **D,** Total percentage of S and G2_M cell cycle stages in REH cells in response to culture conditions as indicated in (A, B) for 72 h (n=3 independent experiments). **E,** Percentage of apoptosis in REH cells in response to culture conditions as indicated in (A, B) and treatment with 150 nM cytarabine (CYT) in RPMI for 72 h (n=3 independent experiments). Data are shown as the mean ± s.e.m. P-values were calculated with a repeated measure one-way ANOVA with Tukey's multiple comparison test (B, E) and two-way ANOVA with Sidak's multiple comparison test (D).

**Supplementary Figure 2: CSFmax culture reduces the consumption rate of metabolites of ALL cells. A,** Extracellular levels of glucose in SEM cells cultured in Plasmax (PL) and CSFmax (CSF) at timepoints 0, 24, 48, and 72 h (n=4 independent cultures). **B,** Percentage of glucose consumption of SEM cells in response to culture conditions as indicated in (A) (n=4 independent cultures). **C,** Extracellular levels of lactate in SEM cells in response to culture conditions as indicated in (A) (n=4 independent cultures). **D,** Percentage of lactate consumption (negative value reflecting secretion) of SEM cells in response to culture conditions as indicated in (A) (n=4 independent cultures). **E, F,** Percentage of consumption of metabolites in SEM cells in cultured in Plasmax (PL) and CSFmax (CSF) for 24 h (E) and 48 h (F) (n=4 independent cultures). Data are shown as the mean ± s.e.m. P-values were calculated with a repeated measure two-way ANOVA with Tukey's multiple comparison test (A, C) and two-way ANOVA with Sidak's multiple comparison test (B, D, E, F).

**Supplementary Figure 3: CSFmax culture reduces the metabolic activity of ALL cells and alters their mitochondrial redox homeostasis. A,** Representative respirometry output profile in SEM cells cultured in Plasmax (PL) and CSFmax (CSF) for 24 h. Arrows represent the time of indicated injection (n=7 independent wells, representative of three independent experiments). **B,** Representative extracellular acidification rate (ECAR) profile in SEM cells in response to culture conditions as indicated in (A). Arrows represent the time of indicated injection. (n=6 independent wells, representative of two independent experiments). **C, D,** Representative respirometry output profile (C) and quantification of maximal respiration (D) in REH cells in response to culture conditions as indicated in (A). Arrows represent the time of indicated injection (n=7 independent wells, representative of three independent experiments). **E, F, G,** Representative extracellular acidification rate (ECAR) profile (E) and quantification of glycolysis (F) and glycolytic capacity (G) in REH cells in response to culture conditions as indicated in (A). Arrows represent the time of indicated injection (n=6 independent wells, representative of two independent experiments). Data are shown as the mean ± s.e.m. P-values were calculated with unpaired t-test (D, F, G).

**Supplementary Figure 4: Intracellular changes in glycolysis and TCA cycle related metabolites in ALL cells cultured in CSFmax. A, B, C, D,** Total intracellular levels of glycolysis related metabolites (A), pyruvate (B), lactate (C), TCA cycle related metabolites (D), reduced glutathione (GSH) (E), and oxidised glutathione (GSSG) (F) in SEM cells cultured in Plasmax (PL) and CSFmax (CSF) for 24 h (n=3 independent cultures). Data are shown as the mean ± s.e.m. P-values were calculated with unpaired t-test (A, B, C, D, E, F).

**Supplementary Figure 5: Autophagy and mitophagy associated genes are upregulated in CNS-derived ALL cells. A,** Normalised expression values of key autophagy associated genes in human SEM and REH cells isolated from the CNS and the spleen of xenografted mice (n=2 independent groups of 5 mice for SEM cells, n=2-3 independent groups of 5 mice for REH cells). Publicly available dataset used: GSE135115. **B,** Logarithmic fold change of autophagy associated genes in human ALL cells isolated from the CSF versus the BM from patient-derived samples with B-ALL at the time of combined BM and CNS relapse (n=3 biological replicates). P-values are indicated next to the bar of each gene. Publicly available dataset used: GSE81519. **C,** Normalised expression values of autophagy associated genes in human ALL cells isolated from the CSF and the BM of patient-derived samples with B-ALL at the time of combined BM and CNS relapse (n=3 biological replicates). Data are shown as the mean ± s.e.m. P-values were calculated with unpaired t-test (A, C).

**Supplementary Figure 6: ALL cells cultured in CSFmax induce and rely on autophagy. A,** Representative western blot showing changes in autophagy and hypoxia-induced markers in SEM cells in response to culture conditions as indicated in (A) (n=2 independent experiments). **B,** Immunoblot analysis showing deletion of BNIP3 in SEM cells cultured in RPMI, in normoxia (N, 21% $O_2$) or hypoxia (H, 1% $O_2$). **C,** Flow cytometry gating strategy to measure leukaemia burden in xenografted mice. **D,** Schematic representation to assess autophagy flux using the GFP-LC3-RFP-LC3ΔG autophagy flux probe. Endogenous ATG4 cleaves the autophagic flux probe into GFP-LC3 (autophagic substrate) and RFP-LC3ΔG (internal control). Reduction in GFP/RFP ratio indicates higher autophagy flux. Schematic diagram is adapted from Kaizuka *et al* [1]. **E,** Schematic diagram of autophagy pathway and the mechanism of action of MRT403 to inhibit autophagy. **F,** Percentage of apoptosis in REH cells cultured in RPMI, Plasmax (PL), and CSFmax (CSF) for 24 h in normoxia (21% $O_2$) with or without 1 µM of MRT403 treatment (n=3 independent experiments). **G,** Percentage of apoptosis in REH cells cultured in RPMI, Plasmax (PL), and CSFmax (CSF) for 24 h in normoxia (21% $O_2$) or hypoxia (1% $O_2$) with or without 50 nM of bafilomycin A1 treatment (n=3 independent experiments). Data are shown as the mean ± s.e.m. P-values were calculated with a repeated measure one-way ANOVA with Tukey's multiple comparison test (F, G).

**Supplementary Figure 7: ATG7 deficient ALL cells increase their uptake of glucose and glutathione in the CNS. A,** Immunoblot analysis of protein expression of ATG7 and autophagy markers in isolated human $CD19^+$ cells retrieved from the BM in xenograft experiment #3 (Exp #3: n=4 mice for ATG7 KO group). **B,** Heatmap showing fold change of intracellular levels of glycolysis, TCA cycle, nucleotides, and other relevant metabolites in human $CD19^+$ SEM cells isolated from the BM and the brain of female NSG immunodeficient mice (n=4 biological replicates). First bar shows fold change of intracellular levels of metabolites between vector control (CTRL) CNS-derived and CTRL BM-derived leukaemic cells. Second bar shows fold change of intracellular levels of metabolites between ATG7 knockout (KO) BM-derived and CTRL BM-derived leukaemic cells. Third bar shows shows fold change of intracellular levels of metabolites between ATG7 KO CNS-derived and CTRL CNS-derived leukaemic cells. **C, D, E,** Intracellular levels of glucose (B), glyceraldehyde-3-phosphate (C), and lactate (D) in human $CD19^+$ SEM cells isolated from the BM and the brain of female NSG immunodeficient mice relative to CTRL BM-derived leukaemic cells (n=4 biological replicates). **F,** Intracellular ATP/ADP ratio in human $CD19^+$ SEM cells isolated from the BM and the brain of female NSG immunodeficient mice relative to CTRL BM-derived leukaemic cells (n=4 biological replicates). Data are shown as the mean ± s.e.m. P-values were calculated with a repeated measure one-way ANOVA with Tukey's multiple comparison test (C, D).

**Supplementary Table 1: Direct comparison of the concentration of key metabolites composing CSFmax with RPMI, Plasmax, and the average CSF metabolome.**

**Supplementary Table 2: Supplements added in RPMI, Plasmax, and CSFmax.**

**Supplementary Table 3: List of guide sequences used to generate knockout cell lines.**

**Supplementary Table 4: List of primary and secondary antibodies used in western blots.**

### Supplemental Materials and Methods

#### Formulation of CSF-like medium (CSFmax)

To produce CSFmax, stock solutions of each component were prepared in individual tubes and stored at -80 °C. To prepare a bottle of CSFmax, a master mix of all stock solutions was prepared and diluted in a buffered solution containing: customised EBSS (without glucose) [2], sodium bicarbonate (Cat. #S5761), phenol red (Cat. #114529), BME vitamin mix (Cat. #B6891), β-mercaptoethanol (Cat. #21985023), and Plasmax™ (Cat. #156371) [3]. CSFmax was supplemented with 100 IU/mL penicillin/streptomycin (Cat. #15140122) and 5% (v/v) delipidated dialysed fetal bovine serum (FBS) (Cat. #A3382001). Dialysed FBS was delipidated using fumed silica precipitation. All components were added in a stericup vacuum filter, and the pH was adjusted to be at 7.2. CSFmax cell culture medium was stored at 4 °C.

#### FBS delipidation

Dialysed FBS was delipidated using fumed silica precipitation. For this, 2 g of fumed silica (Cat. #S5505) were added to 100 mL of serum and were mixed using a magnetic stirrer at room temperature (RT) for 2 h. Following this, the mix was transferred to falcon tubes which were then centrifuged at 10,000 rpm for 30 min at RT. After pelleting the lipids, the supernatant was collected, filter sterilised and stored at -20 °C until further use.

#### Growth curves and proliferation rates

ALL cell lines were seeded at a density of 50,000 cells/mL in a 12-well plate in appropriate culture conditions. Cell counts were obtained at 24 h intervals using the CASY automated cell counter (Roche). The proliferation rate was calculated using the following formula:

Proliferation rate (doublings per day) $=\log_2$(final cell count/initial cell count)/no. of days.

#### Cell cycle analysis

ALL cell lines were cultured at a density of 100,000 cells/mL in appropriate culture conditions for 72 h. Following incubation, cells were fixed in ice- cold 70% ethanol dropwise while vortexed and then incubated at -20 °C for 2 h. Cells were then washed twice in phosphate buffered saline (PBS) solution (Cat. #14190094) and centrifuged at 400 g for 5 min. To ensure that only DNA was stained, cells were treated with 2 µL of RNase A (100µg/mL) (QIAGEN). Cells were stained with 200 µL of propidium iodide (50µg/ml) (PI, Cat. #P1304MP) and were analysed using the FACSVerse flow cytometer (BD Biosciences). Analysis of cell cycle stages was performed with FlowJo V10.8.0. and the geometrical mean intensity of PI was calculated to separate different cell cycle phases.

#### Apoptosis assay

ALL cell lines were cultured at a density of 100,000 cells/mL in appropriate culture conditions and in the presence of treatments for 24 h or 72 h, as indicated. Cells were stained for 15 min at RT with 3 µL 7-aminoactinomycin D (7-AAD, Cat. #559925) and 3 µL Annexin V (APC, Cat #550475) and analysed by flow cytometry using the FaCSVerse flow cytometer (BD Biosciences). Alternatively, cells were stained with 1 µg/mL 4′,6-diamidino-2-phenylindole (DAPI, Cat. #D9542) dissolved in phosphate buffered saline (PBS) solution (Cat. #14190094) and analysed by flow cytometry using the Attune NxT flow cytometer (Invitrogen).

**Autophagy flux analysis**

ALL cell lines expressing the fluorescent probe GFP-LC3-RFP-LC3ΔG were cultured at a density of 100,000 cells/mL in appropriate culture conditions for 24 h. Following incubation, cells were harvested and washed with phosphate buffered saline (PBS) solution (Cat. #14190094). Immediately, cells were analysed using the Attune NxT flow cytometer (Invitrogen). Analysis of the mean fluorescence intensity (MFI) of GFP and RFP were analysed using FlowJo V10.8.0. and the ratio of GFP/RFP was calculated to indicate autophagy flux.

**Mitochondrial ROS measurements**

ALL cell lines were cultured at a density of 100,000 cells/mL in appropriate culture conditions for 24 h and stained with 5 µM MitoSOX Red (Cat. #M36008) in 1 mL phosphate buffered saline (PBS) solution (Cat. #14190094). Cells were incubated for 30 min at 37 °C and analysed by flow cytometry using the FaCSVerse flow cytometer (BD Biosciences).

**Measurement of oxygen consumption rate**

Oxygen consumption rate (OCR) was measured using the XF96 Seahorse Flux analyser (Agilent Technologies). Cells were seeded 24 h prior to the experiment in appropriate culture conditions. The XF96 cell plate was coated with Cell-Tak solution at 0.02 mg/mL in 0.1 M $NaHCO_3$ and pH was adjusted to 7.4. Following a 24 h incubation, ALL cell lines were suspended in XF Assay Medium (Cat. #100965–000) supplemented with 11.1 mM glucose and 2 mM glutamine and seeded at a concentration of 100,000 cells/well in the pre-coated XF96 well plate. Measurement of OCR was performed at baseline and after sequential injections of oligomycin (1 µM), carbonyl cyanide 3-chlorophenylhydrazone (CCCP) (1 µM), and rotenone (1 µM) and antimycin A (1 µM). Maximal respiration was calculated according to manufacturer's instructions.

**Measurement of extracellular acidification rate**

Extracellular acidification rate (ECAR) was measured using the XF96 Seahorse Flux analyser (Agilent Technologies). Cells were seeded 24 h prior to the experiment in appropriate culture conditions. The

XF96 cell plate was coated with Cell-Tak solution at 0.02 mg/mL in 0.1 M $NaHCO_3$ and pH was adjusted to 7.4. Following a 24 h incubation, ALL cell lines were suspended in XF Assay Medium (Cat. #100965–000) supplemented with 2 mM glutamine and seeded at a concentration of 100,000 cells/well in the pre-coated XF96 well plate. Measurement of ECAR was performed at baseline and after sequential injections of glucose (10 mM), oligomycin (1 µM), and 2-deoxyglucose (50 mM). Glycolysis and glycolytic capacity were calculated according to manufacturer's instructions.

**Western blot analysis**

Cells were washed with phosphate buffered saline (PBS) solution (Cat. #14190094) and lysed into RIPA buffer (Cat. #89900) containing cOmplete Protease (Roche) and PhosSTOP phosphatase (Roche) inhibitors. Cell lysates were centrifuged at 16,000 g for 20 min at 4 °C. Protein concentration was quantified by bicinchoninic acid assay (Thermo FIsher Scientific). Lysates were resolved into a 4-12% polyacrylamide gel electrophoresis gel and were transferred to a PVDF membrane. Blocking was performed for 1 h in 2% BSA solution (in Tris-buffered saline, 0.01% Tween). Membranes were incubated overnight with indicated primary antibodies (Supplementary Table 4) at 4 °C. The next day membranes were washed in 2% BSA solution for 1 h with gentle rocking and were incubated with indicated secondary antibodies (Supplementary Table 4) at RT. Bands were detected by ECL reagent (Cat. #RPN2235) and developed using the Odyssey FC imaging system V.5.2.

**Intracellular steady-state metabolic profiling**

To measure intracellular levels of metabolites *in vitro*, SEM cells were cultured at a density of 200,000 cells/mL in appropriate culture conditions for 24 h. After the 24 h incubation period, cell concentration was measured using the CASY automated cell counter (Roche). Whereas, to measure intracellular levels of metabolites *in vivo*, human $CD19^+$ cells were isolated from each organ as described above (see section Cell line xenograft experiments). After ensuring through flow cytometry, that the purification of human $CD19^+$ cells harvested is above 95% in all experimental conditions, metabolites were extracted and analysed.

Cell concentration counts for both *in vitro* and *in vivo* analysis were used to normalise the volume of extraction solvent to achieve a concentration of $1x10^6$ cells/mL in all experimental conditions. Cells were washed twice with ice-cold phosphate buffered saline (PBS) solution (Cat. #14190094) with pulse centrifugation at 12,000 g for 15 sec at 4 °C. Following this, cells were suspended with the appropriate volume ice-cold extraction solvent (50:30:20, v/v/v acetonitrile/methanol/water) and were vortexed for 30 sec and incubated at 4 °C for 5 min. Cells were then centrifuged at 16,000 g for 10 min at 4 °C and the supernatant was transferred to liquid chromatography-mass spectrometry (LC-MS) glass vials. LC-MS glass vials were stored at -80 °C until measurements were performed.

**Extracellular steady-state metabolic profiling**

To measure extracellular levels of metabolites, present in cell culture media, ALL cell lines were cultured at a density of 50,000 cells/mL in appropriate culture conditions. Cells were then centrifuged at 400 g for 5 min at 4 °C, were pelleted, and 50 µL of supernatant (medium) were transferred in LC-MS glass vials containing 1 mL of extraction buffer (50:30:20, v/v/v acetonitrile/methanol/water). LC-MS glass vials were stored at -80 °C until measurements were performed. Notably, after the indicated incubation period and prior to extraction, cell concentration was measured using the CASY automated cell counter (Roche) which was used for quantitative analysis of extracellular levels of metabolites. The peak intensity of each metabolite was normalised to cell number and relative to medium only samples using the following formula:

Exchange rate = Δmetabolite / ((cell number day 0 + cell number day 1)/2)

**Liquid chromatography-mass spectrometry (LC-MS) analysis**

Samples were analysed using a Thermo Ultimate 3000 High pressure liquid chromatography (HPLC) system (Thermo Scientific) coupled to a Q Exactive Orbitrap mass spectrometer (Thermo Scientific). The HPLC system consisted of a ZIC-pHILIC column (SeQuant, 150 × 2.1 mm, 5 µm, Merck KGaA), with a ZIC-pHILIC guard column (SeQuant, 20 × 2.1 mm) and an initial mobile phase of 20% 20 mM ammonium carbonate, pH 9.4, and 80% acetonitrile. For the analysis, 5 µL of metabolite extracts were injected into the system, and metabolites were separated over a 15 min mobile phase gradient decreasing the acetonitrile content to 20%, at a flow rate of 200 µL/min and a column temperature of 45 °C. Metabolites were then detected using the mass spectrometer operating in polarity switching mode. All metabolites were detected across a mass range of 75–1000 m/z at a resolution of 35,000 and greater (at 200 m/z), and with a mass accuracy below 5 ppm. Data were acquired using Thermo Xcalibur software Version 2.11 QF1 Build 3006. The peak intensities of different metabolites were determined using the TraceFinder 4.1 software (Thermo Fischer Scientific) and the Skyline software [4]. Metabolites were identified by accurate mass of the singly charged ion and by known retention times on the pHILIC column. Peak intensities were normalised to cell number and volume measured prior to metabolite extraction.

**Computational analysis**

Publicly available transcriptomic datasets (GSE135115 and GSE81519) were analysed using the R software. The steps followed for the analysis included; quality control and normalisation of the raw data, filtration of low expressed genes to enhance quality of analysis, differential gene expression analysis using the DeSeq2 package. Statistical significance was determined using the Benjamini–Hochberg method. Adjusted $p < 0.05$ and $\log_2 FC \geq 2$ were significant. GSEA (version 4.1.0) analysis was conducted on pre-ranked lists (ranked by pi score calculated by multiplying $Log_2$ FC by –$Log_{10}$ (adjusted *p*-value)).

**Human CD19$^+$ cell isolation and tumour burden assessment**

Mice were sacrificed at experimental endpoint (day 21 post-engraftment) and organs (brain, BM, spleen) were harvested. Brain was harvested and it was transferred to a bijoux vial in ice-cold PBS. Additionally, the inside of the skull was scraped and washed with ice-cold PBS to collect all brain cells and transferred to the same bijoux vial. To process the brain, firstly, it was transferred to a 15 mL falcon tube in ice-cold PBS. Following gentle vortexing of the brain for 30 sec, it was then strained into a 50 mL falcon tube on ice. Any excess cells left in the 15 mL falcon tube were washed with ice-cold PBS and passed through the strainer to bring total volume of cells in PBS to 6 mL. Cells were then added into a 15 mL falcon tube containing 3 mL of Lymphoprep Density Gradient Medium (Cat. #7851). To process the BM and the spleen, they were firstly crushed, washed with ice-cold PBS, and then strained and transferred to a 50 mL falcon tube. Similarly to the brain, total volume of cells in PBS was 6 mL. Cells were then added into a 15 mL falcon tube containing 3 mL of Lymphoprep Density Gradient Medium (Cat. #7851).

To separate the cells through the Lymphoprep Density Gradient Medium (Cat. #7851), tubes were centrifuged at 800 g for 30 mins at 4°C. Following this, separated cells were collected and were washed in ice-cold PBS at 800 g for 10 mins at 4°C. Cells were suspended in 1 mL of ice-cold PBS and concentration of cells was measured using the CASY cell counter. Cells were then stained with anti-mouse CD45-APC (Cat. #561018), anti-human CD45-PE (Cat. #555483), and anti-human CD19- PE-Cy7 (Cat. #557835) for 20 mins at RT in the dark, to distinguish the human cell population. After staining, cells were washed in ice-cold PBS and were suspended in 300 µL of PBS. 50 µL of counting beads were then added, and cells were subjected to flow cytometry analysis to measure the total number of human CD19$^+$ cells. Human CD19$^+$ cells were isolated and counted using CountBright Absolute Counting Beads (Cat. #C36950) according to manufacturer's instructions and were analysed with flow cytometry to measure tumour burden, using FACSVerse flow cytometer (BD Biosciences). The total number of human CD19$^+$ cells was calculated using the following formula:

Absolute count (cells/µL) = [(Number of cell events x Counting beads volume)/(Number of bead events x Cell volume)] x Counting beads concentration (49,500 beads/50 µL volume)

**Data availability**

All data associated with this study are presented in the paper or in the Supplemental Materials.
